# Cellular correlates of cortical thinning throughout the lifespan

**DOI:** 10.1101/585786

**Authors:** D. Vidal-Pineiro, N. Parker, J. Shin, L. French, H. Grydeland, AP. Jackowski, AM. Mowinckel, Y. Patel, Z. Pausova, G. Salum, Ø. Sørensen, KB Walhovd, T. Paus, AM Fjell, for the Alzheimer’s Disease Neuroimaging Initiative, for the Australian Imaging Biomarkers, Lifestyle flagship study of ageing

## Abstract

Cortical thinning occurs throughout the entire life and extends to late-life neurodegeneration, yet the neurobiological substrates are poorly understood. Here, we used a virtual-histology technique and gene expression data from the Allen Human Brain Atlas to compare the regional profiles of longitudinal cortical thinning through life (4004 MRIs) with those of gene expression for several neuronal and non-neuronal cell types. The results were replicated in three independent longitudinal datasets. We found that inter-regional profiles of cortical thinning related to expression profiles for marker genes of CA1 pyramidal cells, astrocytes and microglia during development and in aging. During the two stages of life, the relationships went in opposite directions: greater gene expression related to less thinning in development and *vice versa* in aging. The association between cortical thinning and cell-specific gene expression was also present in mild cognitive impairment and Alzheimer’s Disease. These findings suggest a role of astrocytes and microglia in promoting and supporting neuronal growth and dendritic structures through life that affects cortical thickness during development, aging, and neurodegeneration. Overall, the findings contribute to our understanding of the neurobiology underlying variations in MRI-derived estimates of cortical thinning through life and late-life disease.

## Introduction

The human cerebral cortex undergoes constant remodeling throughout the lifespan. Magnetic resonance imaging (MRI) enables the estimation of structural properties of the human cerebral cortex that can, in turn, be related to cognitive, clinical and demographic data^1,2^, used as biomarkers of neurodegeneration^3,4^ or as high-fidelity phenotypes for genomic studies^5,6^. Yet, we lack data for guiding interpretation of MRI-based phenotypes with regard to the underlying neurobiology^8,9^. Relating MRI-based variations in cortical thickness to specific neurobiological processes is fundamental for our understanding of the normal lifespan trajectories and individual differences in cortical thickness, as well as the nature of cortical abnormalities in brain disorders and diseases such as Alzheimer’s Disease (AD). Here, we approach this question by studying the regional patterns of age-related cortical thinning in relation to those of the expression of genes indexing several neuronal and non-neuronal cell types.

For MRI-based estimates of cortical thickness, the prototypic lifespan-trajectory includes a steep age-related decrease in childhood and adolescence, followed by a mild monotonic thinning from early adulthood and, in many regions, an accelerated thinning from about the seventh decade of life^10,11^. Despite uninterrupted cortical thinning during the lifespan, the underlying neurobiological substrates and processes must be – at least partially - different. Developmental changes in cortical thickness may involve processes such as intracortical myelination, remodeling of dendritic arbours and its components (e.g., dendritic spines), axonal sprouting, and vascularization^12–15^. With advancing age, cortical thinning may, for instance, be associated with neuronal and dendritic shrinkage^16–18^. Cortical thinning in AD could be also influenced by the degree of neuronal loss^19^.

The human cerebral cortex contains many types of cells, which can be classified into neuronal – mostly pyramidal cells and interneurons - and non-neuronal cells - mostly glial cells, namely microglia, astrocytes and oligodendrocytes^19–21^. Histological data suggest that regional variations of cortical thickness may be associated with the number of non-neuronal cells and neuropil volume^23,24^. Yet, *ex vivo* approaches have limitations when it comes to understanding the cellular substrate driving age-related changes in cortical thickness in the general population. Two studies have addressed this issue by combining cross-sectional estimates of cortical thickness during adolescence with quantitative MRI data and gene expression data^24,25^. By combining magnetization transfer and cortical thickness estimates, Whitaker and colleagues^25^ suggested intra-cortical myelination as a primary driver of cortical thinning in the adolescent cortex. On the other side, Shin and colleagues^24^ combined regional profiles of age-related cortical thickness in adolescence and cell-specific gene expression using the Allen Human Brain Atlas (AHBA)^27^. Their results suggested that – across the cerebral cortex – inter-regional variations in cortical thinning are associated with inter-regional variations in the expression of marker genes for CA1 pyramidal cells, astrocytes, and microglia (regions with more expression showing less thinning).

It is, however, unclear to which degree cortical thinning relates to the same gene expression patterns throughout development and aging. The answer to this question has implications for our understanding of the neurobiological foundation for age-related changes in cortical thinning. Thus, the main objective of the study was to shed light on the neurobiological substrates underlying cortical thinning at different stages during the lifespan. Here we ask whether age-related changes in thickness of the human cerebral cortex, assessed *in vivo* with longitudinal MRI, are associated with specific neuronal or non-neuronal cell-types. We approached this question by correlating regional profiles of cortical thinning – during different periods of the lifespan - with regional profiles of cell-specific gene expression. We followed up the results with two supplementary questions: i) Is the cortical thinning – gene expression relationship in young and older adults supported by one or several neurobiological mechanisms? ii) Does the cortical thinning – gene expression relationship extend into to the aging-dementia clinical continuum? Aging is the major risk factor for AD^28^ and there is increasing evidence that the systems that are vulnerable to AD are also susceptible to the effects of age (see ^29–32^). Understanding possible common substrates underlying age-related changes in cortical thickness during development, aging and, dementia can provide valuable understanding of why AD initially targets specific brain networks.

## Results

Briefly, the MRI and gene expression datasets were parsed into 34 cortical regions-of-interest (ROIs) of the left hemisphere^33^. MRI data were drawn from the Center for Lifespan Changes in Brain and Cognition (LCBC) consisting of 4,004 observations from 1,899 cognitively healthy participants (**Table S1**). MRI data were processed through the FreeSurfer pipeline and fitted using generalized additive mixed models (GAMM). Cortical thinning was estimated as the first derivative of the fitted model, and units represent mm of thickness change per year (mm/yr). For gene expression, we used the AHBA, a public resource that includes values of gene expression in multiple regions of the human cerebral cortex. As cell-specific markers, we used genes identified as unique to one of nine cell-types based on single-cell RNA data extracted from the CA1 region and the somatosensory region (S1) in mice^34^. Notably, genes with inconsistent inter-regional profiles of their expression (across donors) and with age-varying profiles were filtered out^24^. As most cortical regions show thinning during the whole lifespan, hereafter we refer to the cortical thinning/thickening estimates by cortical thinning and use the terms “more and less thinning” to denote directionality. Note though that the cortical estimates are measured throughout the thickening/thinning continuum and positive values in the derivative reflect thickening while negative values denote thinning.

In addition to the main results, we created an interactive resource using the “shiny” R package to allow an interactive visualization of both the trajectories of cortical thickness and thinning throughout the lifespan, and the inter-regional profiles of cortical thinning at any age (**<u>supporting information</u>**).

### Lifespan trajectories of cortical thickness and cortical thinning

For most regions, the lifespan trajectories of cortical thickness displayed the prototypic pattern characterized by a steep decline in MRI-based estimates of thickness during childhood and adolescence, a mild monotonic thinning from the early adulthood onwards, and a trend towards accelerated thinning in older adulthood (see **Figure 1)**. See an interactive visualization of cortical thickness and thinning trajectories for each region in **<u>supporting information</u>**.

**Figure 1.**
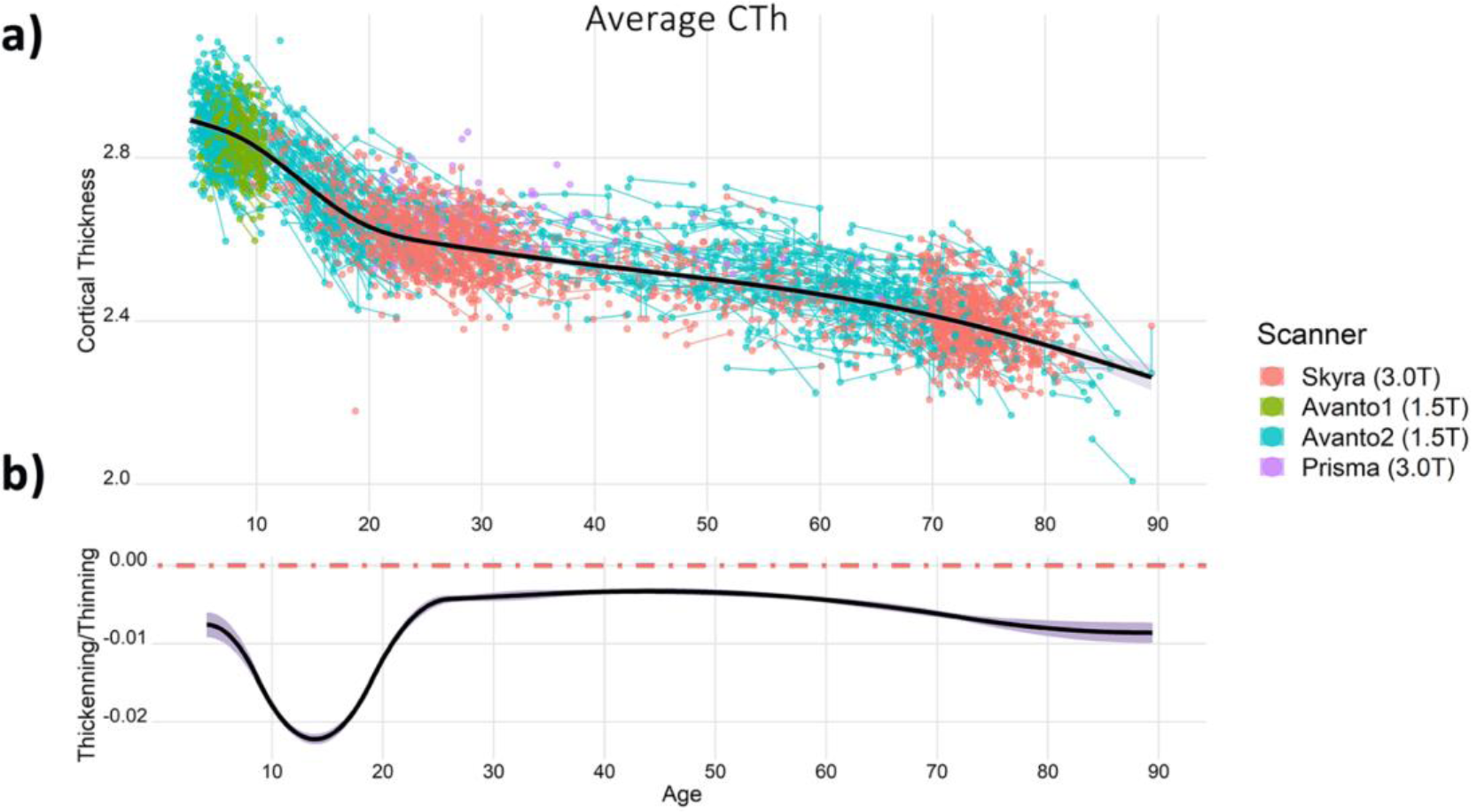
Trajectories of weighted-average cortical thickness. The upper and lower plots exhibit the trajectories of cortical thickness and cortical thinning during the lifespan, respectively. Cortical thickness fitting (black line) overlies a spaghetti plot that displays each observation (dots), participant (thin lines) and, scanner (color). The y-axis units represent mm and Δmm/year for the thickness and thinning plots, respectively. The dotted red line in the cortical thinning graph represents 0 change, negative and positive values represent thinning and thickening, respectively.

### Lifespan Virtual Histology. Cell type – cortical thinning relationship throughout the lifespan

Next, we used the virtual histology approach to assess the relationship between cortical thinning across the lifespan and cell-specific gene expression across the 34 cortical regions. In this analysis, we used solely genes with consistent interregional profiles across individual donors and donor age groups. As shown in **Figure 2a**, we observed that the average correlation coefficient for the different cell-types varied throughout the lifespan. The average correlation coefficients differed from the empirical null distributions for the following cell-types and age periods: astrocytes (5 - 11 years, 35 – 41 years, 61 - 67 years), microglia (5 - 13 years, 65 - 71 years) and CA1 pyramidal cells (5 - 9 years, 63 - 79 years). All results were corrected for both the within (Bonferroni) and between (False Discovery Rate, FDR) cell-type multiple comparisons. Note that the first derivative denotes the degree of thinning so a positive correlation implies that the higher the cell-specific gene expression values are, the less steep the cortical thinning is, and *vice versa*. The average correlation coefficients in childhood/adolescence (for astrocytes, microglia and, CA1 pyramidal cells), and during middle-age (for astrocytes) were positive: thus, cortical regions with higher cell-specific expression for astrocytes, microglia, and CA1 pyramidal cells displayed *less* pronounced thinning. During older age, the gene expression – thinning correlations were negative: thus, cortical regions with higher cell-specific expression for astrocytes, microglia and, CA1 pyramidal cells displayed *more* pronounced age-related cortical thinning. The results were robust to variations of different fitting parameters (**Figure S1, SI Methods, and Results**). The complete statistics are available in **<u>supporting information</u>**. Next, we replicated the main findings in three independent datasets that corresponded to the age periods with significant results (i.e. childhood/adolescence, middle-age and older age periods). As seen in **Figure 2b**, we replicated all expression – thinning correlations observed in the discovery sample: positive correlations for astrocytes, microglia and CA1 pyramidal cells in childhood and adolescence, positive correlation for astrocytes in middle-age, and negative correlations for astrocytes, microglia and CA1 pyramidal cells in older age. Note that significance was only tested for the cell types that were significant in the main (discovery) results. See **SI Methods and Results** for a detailed description of the replication results and the samples used.

**Figure 2.**
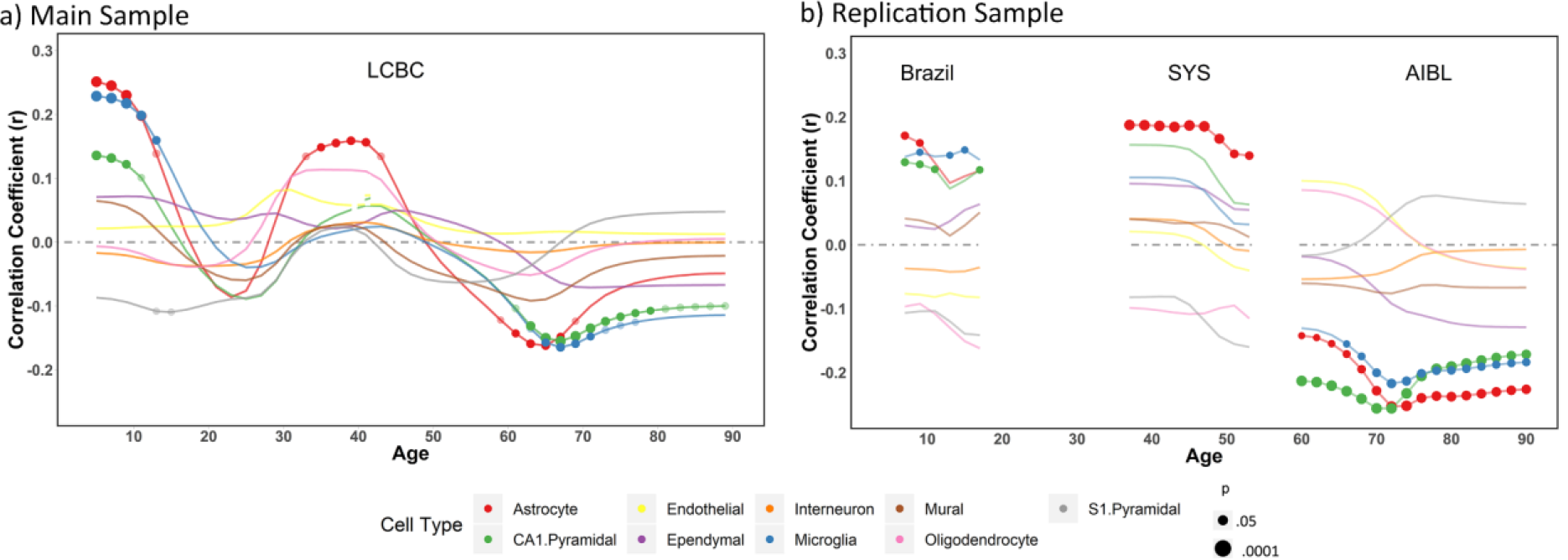
Virtual Histology through the lifespan. Correlation coefficients between the cortical thinning profile through the lifespan and mean regional profiles of gene expression levels in each of the 9 cell types. a) Thinning profile obtained from the LCBC dataset. (n = 4,004 observations). b) Thinning profile for the replication datasets. SYS replication dataset visualization is trimmed after 55 years to avoid overlap with the AIBL sample (see **SI Methods** for more information). Three different sample cohorts were used corresponding to the age-ranges where significant associations between the thinning profile and cell-specific gene expression were found. We used the Brazil High-Risk Cohort to encompass school years, the SYS dataset for middle-aged participants, and the AIBL data for older adults (n = 1,174, 548 and, 739 observations, respectively). Significance was tested only for significant thinning – expression correlations in the main sample; thus expression – thinning trajectories of the additional cell-types are included for visual purposes only. For both plots, the x-axis indicates age while the y-axis indicates the correlation of thinning with cell type-specific gene expression as derived from the Allen Human Brain Atlas. Values above 0 represent a relationship of gene expression profiles with reduced thinning - or thickening - while values below 0 represent a relationship with steeper cortical thinning. Circles indicate a significant relationship (p < 0.05, permutation inference n = 10,000 iterations) after Bonferroni correction for multiple comparisons through the lifespan (semi-transparent circles) and after additionally applying FDR-adjustment for testing multiple cell-types (opaque circles). The size of the circle represents significance. See the cortical thinning profiles and the expression – thinning correlation results in **Figure 3** and **Table S2**. Post-hoc we tested whether this relationship was evident only for specific subclasses of astrocytes, microglia and CA1 Pyramidal cells. In CA1 pyramidal cells, only expression for the CA1Pyr2 subclass was associated with young and old cortical thinning profiles (**SI Methods and Results; Table S3**).

**Figure 3.**
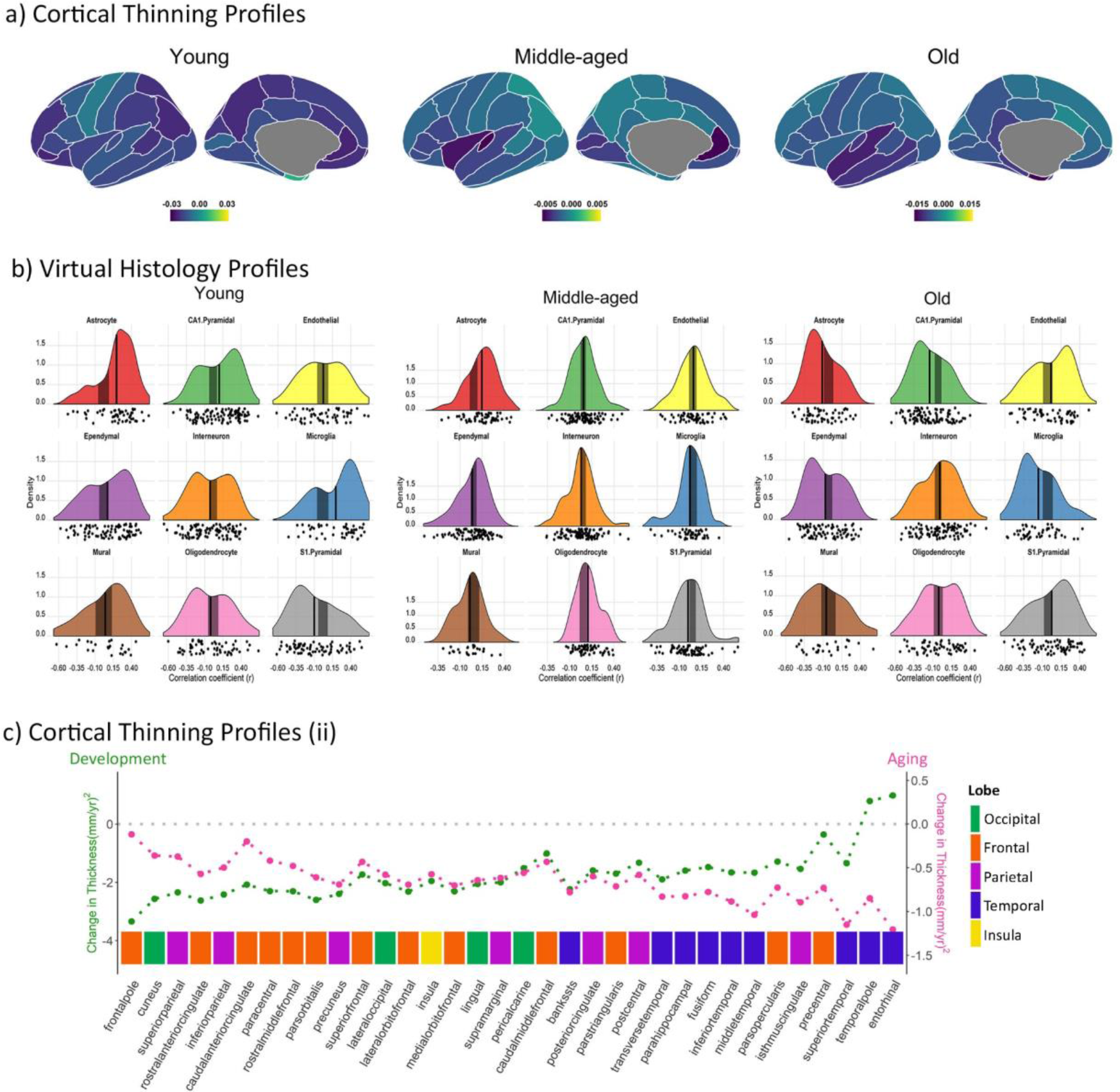
Inter-regional profiles of cortical thinning and Virtual Histology at the ages of interest. a) Inter-regional patterns of cortical thinning for the age-ranges of interest (i.e. age-periods with significant cell-specific expression-thinning associations), as observed in the LCBC dataset. b) Virtual Histology results (expression – thinning association) for the age-ranges of interest, as observed in the LCBC dataset. Each plot shows the distribution of the expression - thinning correlation coefficients for genes in each cell-type group as a density function and as a cloud of dots. The x-axes indicate the coefficients of correlation between the thinning and the expression profiles. The y-axes indicate the estimated probability density for the correlation coefficients; the vertical black line indicates the average correlation coefficient across all genes while the shaded gray box indicates the 95% limits of the empirical null distribution (unadjusted). From left to right: childhood (5 – 9 years), middle-age (35 -41 years), and old age (63-67 years). See also **Table S2**. c) Inter-regional patterns of cortical thinning during childhood (5 – 9 years) and at older age (63-67 years). The y-axes denote cortical thinning while the x-axis represents each cortical region of the left hemisphere. Regions are also categorized by lobe. Note the inverse inter-regional profile of cortical thinning at both ends of the lifespan.

### Virtual Histology. Cell type – cortical thinning relationship in AD

Next, we tested the relationship between cell-specific gene expression and the patterns of cortical thickness decline related to AD and Mild Cognitive Impairment (MCI). We used the ADNI and AIBL as discovery and replication datasets. We fitted cortical thickness using general linear mixed models (*lme4* R-package), with age, sex and, clinical diagnostic (AD, MCI and cognitively healthy controls [HC]) as fixed effects and participant identifier as a random intercept. The results showed that the different patterns of cortical thinning associated with MCI and AD (AD vs. HC, AD vs. MCI and MCI vs. HC) were all significantly associated with the interregional patterns of gene expression for CA1 pyramidal cells, microglia and, astrocytes (see **Figure 4**). Regions with more expression of these markers showed more pronounced thinning. These results were replicated in the AIBL sample. See **Table S4** for complete stats.

**Figure 4.**
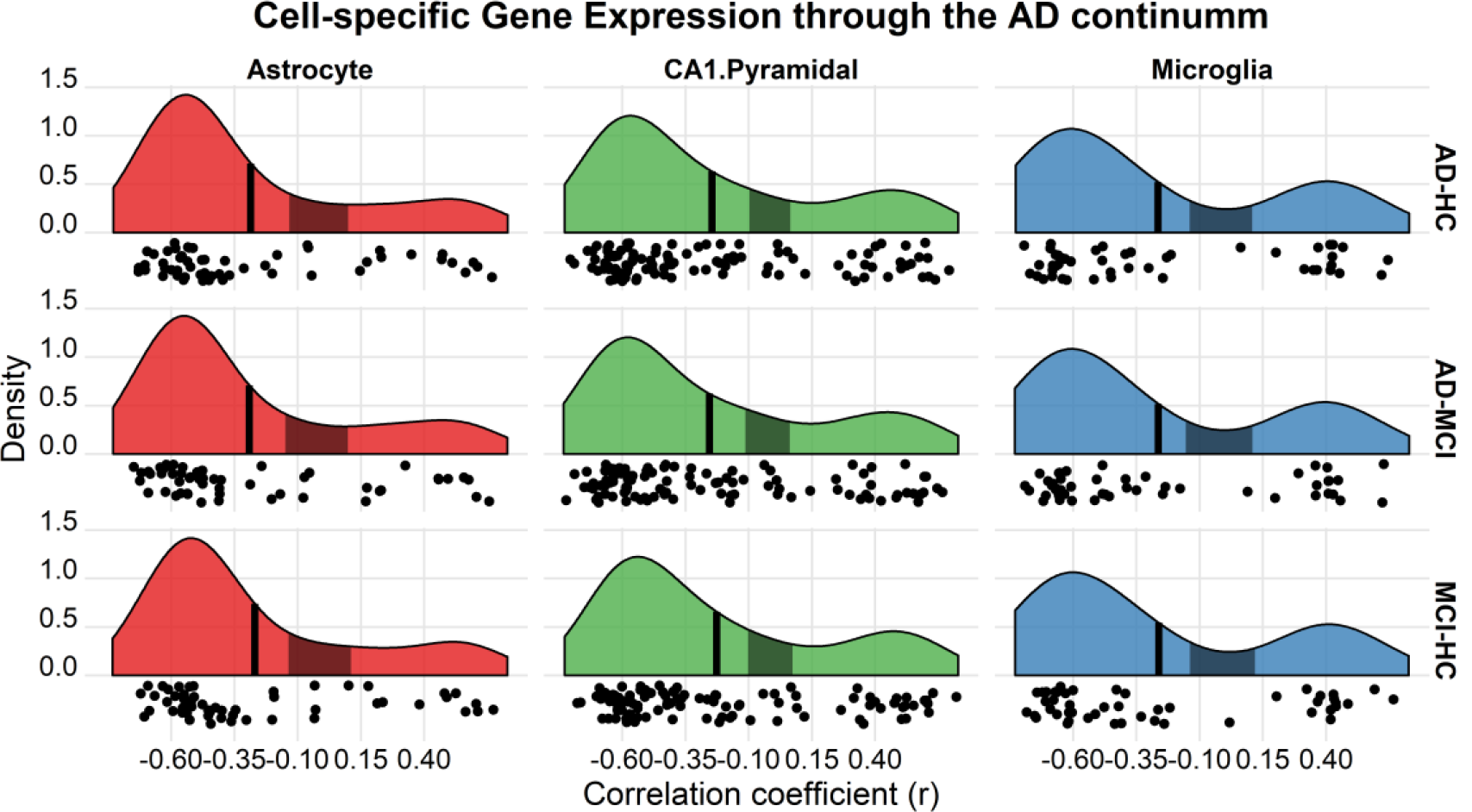
Virtual Histology in AD patterns of cortical thinning. Relationship between cell-specific gene expression profiles and the cortical decline in AD (AD vs. HC; AD vs. MCI; MCI vs. AD). Only results for the CA1 pyramidal, microglia and, astrocyte cell types are displayed. See **Table S4** for complete stats. Each plot shows the distribution of the expression – thinning correlation coefficients for genes in each cell-type group. x-axes = coefficients of correlation between the thinning and the expression profiles. y-axes = estimated probability density for the correlation coefficients. Vertical black line denotes the average correlation coefficient across all genes while the shaded gray box indicates the 95% limits of the empirical null distribution.

### A cortical thickness variation mode as a common mechanism at young and old age as well as in AD

Finally, we tested whether the gene expression – cortical thinning relationships at both young and older age are driven by common or different neurobiological mechanisms. For this, we decomposed inter-individual variability in cortical thickness using an Independent Component Analysis (ICA) approach^35,36^. This ICA decomposition approach results in a set of different modes of variation in cortical thickness that can be attributed to plausible, unique, underlying biological mechanisms. Evidence in favor of common mechanisms for the gene – thinning correlations in young and older age should meet the following conditions: i) Evidence of a cortical thickness independent component (IC) with a strong relationship with age. ii) This IC must follow an inverted U-shape lifespan trajectory as the gene expression – cortical thinning relationship at young and old age have opposite directions and; iii) have to relate to the gene expression profiles for CA1 pyramidal, astrocytes, and microglia cell types.

The results showed three IC components with a relationship with age as assessed with GAMM (r^2^ > .15). See **SI Methods** for details. IC1 represented a dominant whole-brain mode of variation; i.e. showing a topological pattern comparable to the mean cortical thickness (r = .45) and a similar lifespan trajectory as the one exhibited by mean cortical thickness, with the steepest decline in late adolescence (highest rate of decline ≈18 years of age). A post-hoc analysis (**SI Methods**) suggested that the IC1 cortical loadings are topologically associated with age-related changes in the T1w/T2w index - a putative index of myelination - during this adolescent period (r = .42, p = .014, n = 34 ROIs based on Desikan-Killiany atlas) (**Fig 5a**). This result suggests that some changes in cortical thinning – especially in late adolescence – may be best represented by mean cortical thinning and thus cannot be well-captured by interregional differences in cortical thinning. IC2 component largely captured differences of cortical thickness across scanners as the age effects largely overlap with the density distribution of the different scanners (**Fig 5b**). The utility of the ICA approach for denoising has been discussed elsewhere^37^. Finally, the IC3 component satisfied the predefined conditions. It was strongly related to age (r^2^ = .44) and showed an inverted U-shape lifespan trajectory with a steep increase during childhood and a strong decline in old adulthood. Also, the cortical loadings of IC3 were related to the interregional patterns of gene expression for CA1 pyramidal, astrocytes and microglia cell-types (all p_fdr_ < .01; the remaining cell type panels were not associated with IC3 p_fdr_ > .05). We ran a post-hoc analysis to test whether genes that co-expressed with cell-specific genes with high fidelity to IC3 were enriched for AD-related genes. The protocol is thoroughly described by Sliz and colleagues^38^ and in **SI Methods**. The results of the co-expression, enrichment analysis showed that the set of genes that co-expressed with high IC3 fidelity, microglia-specific, genes were significantly enriched by genes associated with AD (p_fdr_ = .01; **Fig 5c**). CA1 pyramidal (p_fdr_ = .09) and astrocytes (p_FDR_ = .24) co-expressed genes were not enriched for AD genes. Altogether, the results suggest the existence of 1) common mechanisms behind the relationship between cortical thinning and specific cell types at both young and old age that 2) are also disproportionately co-expressed with AD-related genes (only for microglia-specific genes). 3) The presence of different modes of cortical thickness variation during the lifespan that can integrate disparate findings in the literature; e.g. myelination vs. dendritic arborization.

**Figure 5.**
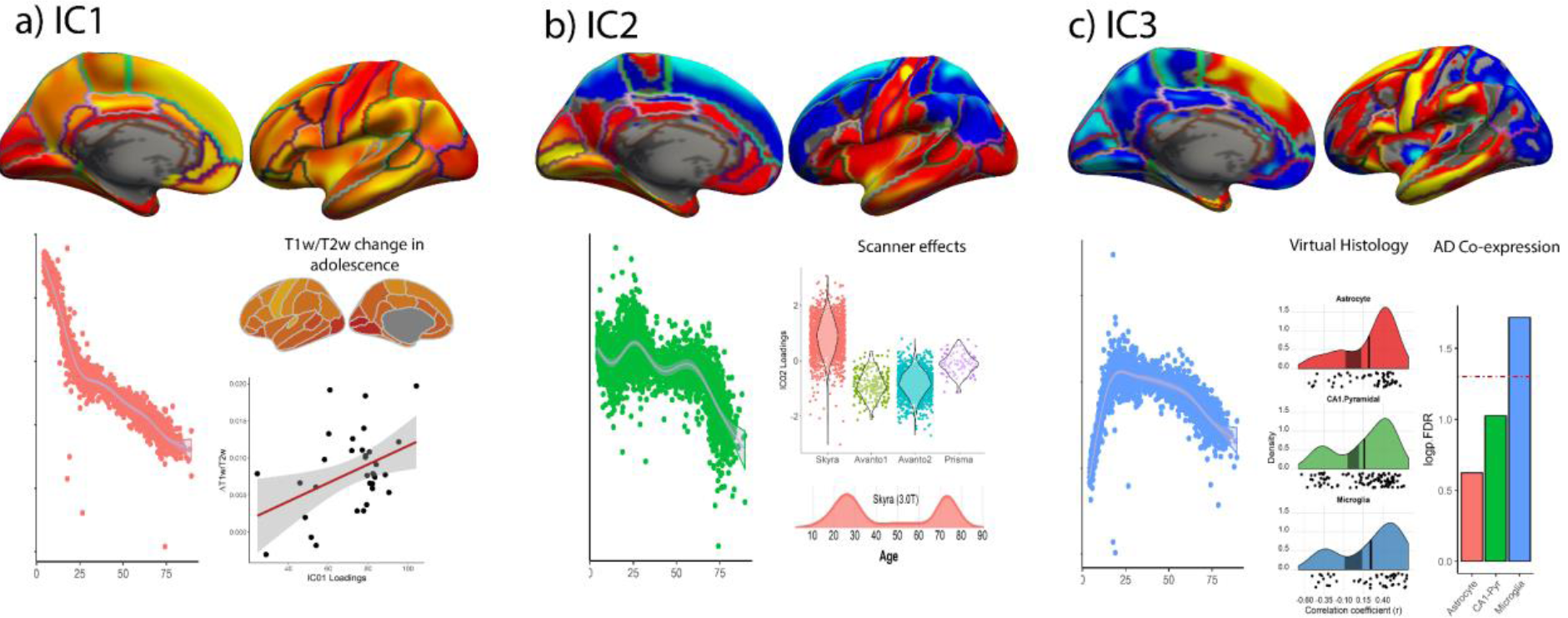
Independent components of cortical thickness. Cortical thickness ICs with practical significance with age (r^2^ = .15; arbitrarily thresholded at Z = 4 for visualization). a) Loadings of IC1 through the lifespan. Increase (red-yellow scale) in T1w/T2w ratio during adolescence (mean derivatives between age 15 and age 21). Relationship between T1w/T2w increase during adolescence and IC1 loadings. b) Loadings of IC2 across the lifespan. Loadings of IC2 grouped by scanner. Density function of the Skyra scanner across the lifespan. c) Loadings of IC3 across the lifespan. Relationship between interregional weights of IC3 and cell-specific gene expression for microglia, CA1 pyramidal and, astrocytes. Relationship between AD genes and high-fidelity IC3 cell-specific genes.

## Discussion

In this study, we showed that inter-regional profiles of cortical thinning estimated from MRI relate to expression profiles for marker genes of CA1 pyramidal cells, astrocytes and microglia. The relationships were particularly evident during development and aging - higher expression of cell-specific genes was associated with less thinning at younger age (< 13 years of age) and steeper thinning in older age (59 years of age). Similarly, gene expression for CA1 pyramidal cells, astrocytes and microglia were also related to the patterns of cortical thinning both in MCI and AD indicating a continuity throughout development, aging, and disease. The implications of the results are discussed below.

We found that cell type expression - thinning patterns during development are mirrored later in life. This overarching pattern fits with the view that thinning during developmental and old age periods closely resemble a topography based on common genetic influences^8^. As cortical thinning and thus the expression - thinning correlations change over the lifespan, the associations can be understood at least from three different angles: 1) cortical thinning reflects a direct loss of cell numbers or – more likely – cell volume; 2) cell-specific processes – e.g. apoptosis or metabolic demands –mediate indirectly the relationship between cell-type and thinning of the cerebral cortex; and 3) an interaction between developmental (or neurodegenerative) processes and MRI acquisition parameters or analytical features – e.g. shifts in T1/T2 boundary delineation may occur due to myelination^11^.

The association of astrocytes, microglia and CA1 pyramidal cells with less cortical thinning during development replicate our earlier findings in an independent, cross-sectional adolescent sample^24^. The involvement of CA1 pyramidal cells with less cortical thinning is in agreement with the notion that axonal sprouting and remodeling of dendritic arbor are associated with relative thickening of the developing cerebral cortex^24^. CA1 pyramidal genes are highly enriched for terms such as “regulation of dendrite extension” in contrast with S1 pyramidal genes, enriched by terms associated with potassium transport and activity^24^. A plausible theory is that the thinning profiles in childhood and early adolescence are partly dependent on the regional dynamics and regional differences in dendritic growth^39,40^. Note that these findings are not in opposition – but rather complementary - with evidence that myelin is an important process behind cortical thinning during development^23,25^. See *considerations* for an extended discussion.

In aging, regions with high CA1 pyramidal expression coincide with those undergoing steeper thinning. This observation fits with both existing histological evidence and current predictions that link dendritic shrinkage with reduced cortical thickness in older brains^16–18,41^. Although neuronal numbers seem preserved in aging^20,41^, recent evidence suggests that specific neural subpopulations numbers – namely neurons with large cell bodies (and possibly larger dendritic arbour) – might decrease in aged brains^42^. Again, S1 pyramidal expression was unrelated to thinning with advancing age, which is consistent with evidence that neuron-specific changes of gene expression with age are dependent on cellular identity^42,43^. A promising direction requires linking variations of neuronal properties^45^ – including regional variations in dendritic and soma size - with MRI estimates of cortical thinning. As a proof of principle, we found that for CA1 pyramidal cells, only expression for the *CA1Pyr2* subclass was associated with the thinning profiles (**SI Methods and Results**). Expression of *CA1Pyr2*-specific genes is associated with mitochondrial function, which correlates with the length of projections in cortical neurons^34^.

Expression of genes specific to astrocytes and microglia was associated with cortical thinning in both younger and older ages. Despite modest changes in the cell numbers^20,45^ and cell morphology throughout the lifespan^46,47^, it is more likely that the relationship between cortical thinning and these glial subpopulations relates to interactions with other neural components such as astrocytic metabolic processes and signaling pathways for microglia. Astrocytes distribute energy substrates – mostly lactate – to active neurons^48,49^, sustaining glutamatergic neurotransmission, synaptic plasticity, and thus long-term memory formation^50^. Microglia engage in crosstalk with neurons and astrocytes, can modulate synaptic homeostasis, remodel synaptic structures (e.g., dendritic spines), and is closely involved in the maintenance of neuronal structure and function. Likewise, both cell-types are up-regulated homogenously in the cerebral cortex with advancing age^42,43^, leaning towards a reactive phenotype for astrocytes and a shift to a pro-inflammatory profile for microglia^51,52^. Altogether, we speculate that in childhood and adolescence, astrocytic and microglia relate to regional preservation of cortical thickness by promoting and supporting neuronal development, such as dendritic arborization and synaptic remodeling. In aging, failure or dysregulation of upstream microglia and astrocyte cellular processes might lead to dendritic and synaptic changes and subsequent loss of cortical thickness.

Expression – thinning associations during the lifespan were reproduced in AD. That is, gene expression for microglia, astrocytes and, CA1 pyramidal cells related to the characteristic pattern of cortical thinning in AD. Synapse loss is a key feature of AD^53,54^ while astrocytes and microglia play various roles in the AD cellular and molecular cascade^30^. Astrocytes produce ApoliproteinE^55^ and have a possible role in glucose hypometabolism^56^ and together with microglia they adopt a reactive profile and play a fundamental role in the inflammatory changes in the brain^51,52^. While the results likely reflect several molecular mechanisms, they hint to a last-in, first-out lifespan system with heightened vulnerability to AD. One may speculate, the system is characterized by neural bodies with higher energetic demands, more complex synaptic and dendritic patterns and higher plasticity. In any case, the continuous relationship between cell-specific gene expression and cortical thinning through development, aging and disease points both towards the notion of system’s vulnerability and that – to some extent - differences in AD (vs. MCI and cognitively healty older adults) can be better clarified by understanding age-related changes in healthy aging.

### Considerations

Two features of the study require special consideration, namely the source of variability and the correlational nature of the analyses. Correlational tests are unavoidable when exploring the underlying mechanisms of cortical thinning *in vivo*, yet longitudinal comparisons (i.e. individual rates of thinning) will approximate better the dynamic relationship of thinning and its neurobiological foundations across the lifespan.

The main measure of the study is the interregional profile of cortical thinning at different ages. While it represents a robust and stable measure, the interregional thinning profile is unable to provide information neither at an individual level nor across time; hence, a portion of the variability remains unexplored. The relative disparity amongst studies might reside on different characterizations of cortical thinning variability, especially during development. In particular, MRI estimates of myelin content have repeatedly been associated with steeper cortical thinning in adolescents^23,25^. Here, lack of findings may relate to the nature of the interregional profiles, which represent snapshots in time and, hence, are determined by the differential sum of processes that modulate cortical thickness in each area. In other words, interregional comparisons cannot evaluate mean (whole-brain) cortical thinning. Further, the cellular correlates of cortical thinning during childhood in the present study relate to *spared* thinning, which points out that these mechanisms are supplementary to those that drive cortical thinning during adolescence. The ICA analysis tentatively integrates both findings. On one side, we found a main component of cortical thickness variability that displayed topological and lifespan trajectories – such as a steep decline in adolescence - that evoke those of mean cortical thickness. This component related to T1w/T2w signal increase during adolescence, a putative index of myelination^57^. One can speculate that this pattern is concealed in the interregional approach due to its similarity to mean cortical thinning^25^. On the other side, a second component linked both the expression – thinning relationships during development and aging and further extend it to AD. As interregional variability in cortical thinning is quite substantial – with some regions showing significant thickening during childhood – the results pinpoint to the presence of complementary processes in development that require further understanding. Altogether, the results suggest that, at least in early phases of life, diverging mechanisms promote at the same time thinning and thickening of the cerebral cortex^41^.

## Conclusion

Understanding the biological substrate underlying MRI-based estimates of cortical thinning during different stages of the lifespan remains a key issue in neuroscience given the ever-increasing use of neuroimaging and the association of variations in cortical thickness with neurodevelopmental and neurodegenerative pathology. We found that cortical thinning was associated with the expression of genes specific to CA1 pyramidal cells, astrocytes, and microglia in both development and in aging. This expression – thinning relationship was conserved in AD. The study informs of a common biological substrate that affects cortical thickness and links development, aging and systems vulnerability to AD. We speculate that microglia and astrocytic functions are essential for promoting and supporting neuronal growth and development in childhood and thus relative regional preservation of thickness in childhood. Failure of these upstream processes might lead to dendritic changes and subsequent cortical thinning in aging.

## Material and methods

### MRI sample

A total of 4,004 scans from 1,899 healthy participants (1,135 females) ranged between 4.1 – 89.4 years of age (mean visit age = 37.4 [SD = 25.3]) (**Figure 1; Table S1**) were drawn from six Norwegian studies coordinated by the Center for LCBC: The Norwegian Mother and Child Cohort Neurocognitive Study^58^; Neurocognitive Development^59^; Cognition and Plasticity Through the Lifespan^60^; Constructive Memory^61^; Method of Loci^62^; and Neurocognitive plasticity^63^. More than one scan was available for 1,017 participants; the mean number of scans per participant was 2.1 (range 1-6) spanning up to 11.1 years after the initial scan. The density of sampling was higher in childhood/adolescence and during the 7^th^ and 8^th^ decade of life. See **Figure S2** and **Table S1** for visual representation and descriptive information.

In all studies, participants were screened through health and neuropsychological assessments. Common exclusion criteria consisted of evidence of neurologic or psychiatric disorders, learning disabilities or current use of medicines known to affect the nervous system. In addition, participants were excluded from the current study based on the following criteria: lack of a valid structural scan (i.e. excessive movement, acquisition or surface reconstruction errors), score <25 on the Mini-Mental State Examination (MMSE)^63^, score of ≥20 on the Beck Depression Inventory (BDI)^65^, and age > 90 due to low sampling density after this age. All the studies were approved by a Norwegian Regional Committee for Medical and Health Research Ethics. See **SI Methods** for further details on samples.

For replication, we used three additional samples that encompassed the age periods in which we observed associations between regional patterns of cortical thinning and cell-specific gene expression. We used the Brazil High-Risk Cohort^66^ to replicate findings during childhood/adolescence. For replication of results in middle-age, we used the cohort of parents of the Saguenay Youth Study (SYS)^67^. For older adults, the replication sample was collected by the AIBL study group. AIBL study methodology has been reported previously^68^. See **Figure S2, Table S5** and, **SI Methods** for the replication sample description.

We used the ADNI database (adni.loni.usc.edu)^69^ (discovery) and AIBL^68^ (replication) samples to estimate the patterns of cortical thinning in AD. ADNI was launched in 2003 as a public-private partnership, led by PI MW Weiner. The primary goal of ADNI has been to test whether serial MRI, other biological markers, and clinical and neuropsychological assessments can measure the progression of MCI and early AD. The *Australian Imaging, Biomarker & Lifestyle Flagship Study of Ageing (AIBL)* took place in Australia, beginning in 2006, with the aim to study which biomarkers, cognitive characteristics, and health and lifestyle factors determine the development of AD. See **Table S6** and **SI Methods** for further description

### MRI acquisition and preprocessing

LCBC image data were collected using four different scanners. Three scanners (*Avanto2 [1*.*5 T], Prisma [3*.*0 T] and Skyra [3*.*0 T];* Siemens Medical Solution) were located at Oslo University Hospital Rikshospitalet while the remaining (*Avanto1 [1*.*5 T])* was placed at St. Olav’s University Hospital in Trondheim. For 285 participants, data were available for two or more MRI machines from the same day (n = 285). For each participant and visit, we obtained a T1w magnetization prepared rapid gradient echo (MPRAGE) sequence. See **SI Methods** for MRI acquisition parameters in both the LCBC and the replication datasets.

Data were processed on the Colossus processing cluster, University of Oslo. We used the longitudinal FreeSurfer v.6.0. stream^70^ for cortical reconstruction of the structural T1w data (http://surfer.nmr.mgh.harvard.edu/fswiki)^71–73^. See the pipeline description in **SI Methods**. See **SI Methods** for differences in the MRI processing pipeline in the replication datasets.

### MRI analysis. Estimation of cortical thinning through the lifespan

After the surface reconstruction, MRI-based estimates of cortical thickness were extracted for the 34 left hemisphere ROIs corresponding to the Desikan-Killiany atlas^33^. We ran an analysis to estimate each region’s cortical thinning - or thickening - at any given point during the lifespan. First, for each of the 34 cortical regions, thickness data were fitted to age using GAMM with “mgcv” R package^74^. The GAMM fitting technique represents a flexible routine that allows nonparametric fitting with relaxed assumptions about the relationship between thickness and age^8^. The technique is well suited for fitting nonlinear relationships through local smoothing effects, independent of any predefined model, and robust to age-range selections and distant data points^75^.

All GAMM models included sex and scanner as covariates and participant identifiers as random effects. We specified cubic splines as smooth terms^74^ and limited to k = 6 the knots of the spline, a somewhat conservative option to minimize the risk of overfitting and to increase the stability of the derivatives. Next, we excluded those observations ± 7 SD above or below the cortical thickness trajectory (mean = 3.2 [1.8 SD] observations excluded per region). After outlier removal, cortical thickness data were re-fitted by age with GAMM (separately in each region).

Finally, we considered the GAMM derivatives to obtain the trajectories of cortical thinning throughout the lifespan. For each region, we obtained the derivative – and its confidence intervals - based on a finite differences approach as implemented in the “*schoenenberg”* R package (https://github.com/gavinsimpson/schoenberg). Positive values in the derivative are interpreted as steeper thickening while negative values are interpreted as steeper thinning at any given age. For each cortical region, we then extracted the derivative values at discrete ages (from 5 to 89 years; every 2 years) resulting in n = 43 cortical thinning profiles spanning from childhood to old adulthood.

### Gene expression analysis. Estimation of cell-specific gene expression across the cortical surface

We used the scripts provided by Shin and colleagues^24^ to obtain gene expression profiles across the left hemisphere. Note that only genes with inter-regional profiles consistent across the six donors (AHBA) and across two datasets (AHBA and the BrainSpan Atlas) were used. See **SI methods** and elsewhere and elsewhere ^24^ for details.

#### Gene expression profile

Gene expression data were obtained from the AHBA, based on postmortem human brains, providing comprehensive coverage of the normal adult brain (http://www.brain-map.org)^27^. Isolated RNA was hybridized to custom 64K Agilent microarrays (58,692 probes) by the Allen Institute. Gene expression data for the left hemisphere were available for six donors.

As described previously^76^, gene-expression data from the AHBA were mapped to the Desikan-Killiany atlas in FreeSurfer space yielding up to 1,269 labeled samples per brain inside or close to a FreeSurfer cortical region^33^. The expression values of the mapped samples were mean averaged across microarray probes to provide a single expression value for each gene for a given sample. Median averages were used to summarize expression values within each ROI and donor, which was followed by the median average across the six donors. This yielded a single value for each region representing the median profile for a given gene across the 34 cortical regions.

#### Panels of Cell-Specific Marker Genes

Lists of genes expressed in specific cell types were obtained from Zeisel and colleagues^34^. In their study, single-cell transcriptomes were obtained for 3,005 cells from the somatosensory cortex (S1) and the CA1 hippocampus region of mice. Gene expression was then biclustered into nine classes that contained each over 100 genes. We converted mouse genes to human gene symbols as implemented in the *“homologene”* R package^77^. The resulting classes of cell types and the number of marker genes kept after consistent genes were filtered are: S1 pyramidal neurons (n = 73 human gene symbols), CA1 pyramidal neurons (n = 103), interneurons (n = 100), astrocytes (n = 54), microglia (n = 48), oligodendrocytes (n = 60), ependymal (n = 84), endothelial (n = 57), and mural (pericytes and vascular smooth muscle cells; n = 25).

### Higher-level statistical analysis

All the in-house procedures were implemented in R (www.r-project.org). Figures were created using ggplot, FreeSurfer viewers and the ggseg package^78,79^.

#### Correlation between lifespan thinning and cell-specific gene expression profiles

In the main analysis, we assessed the relationship between the inter-regional profiles of cortical thinning – obtained from the GAMM derivative sampled between 5 and 89 years - and the inter-regional profiles of gene expression associated with specific cell types. For each cortical profile and marker gene, we computed the Pearson correlation coefficients between the thinning profile and the median inter-regional profile of gene expression levels for each marker gene; yielding a gene-specific measure of expression - thinning correlation.

We used a resampling-based approach to test the association between cell-type expression and cortical thinning profiles^24^. The average expression - thinning correlation for each group of cell-specific genes served as the test statistic (i.e. mean correlation between cortical phenotype and cell-specific gene expression). In addition, we controlled for multiple comparisons, accounting both for the number of cortical thinning profiles (i.e. age points n = 43) and for the different cell-types (n = 9).

For each cell-type panel, we obtained the empirical null distribution of the test statistic as follows. 1) From all the (n = 2,511) consistent genes, we randomly selected the same number of genes as that of the genes in the cell-type group under consideration. 2) For each thinning profile (n = 43 sampled ages), we averaged the expression - thinning correlation coefficients of all the selected genes. Thus, for each random sample, we obtained the mean correlation coefficients at each age. 3) Of the n = 43 coefficients, we selected the one with absolute maximum expression - thinning correlation. This step is essential to introduce a Bonferroni-like correction for multiple comparisons at a within cell-type level while accounting for the non-independency of the thinning estimates (e.g. the cortical thinning profiles at age 60 and 62 are not independent observations). The null distribution was estimated by repeating the previous three steps for 10,000 iterations. A two-sided p-value was computed as the proportion of average thinning – expression correlations whose values exceeded the average for the genes in the original cell-type group. We rejected the null hypothesis at p = .05 FDR-adjusted significance thus correcting for multiple testing of the nine cell-type groups.

The same procedure was run for the three replication-samples with the following variations. The number of within cell-type comparisons was defined by the specific age range of each replication sample (n = 6, 15 and 17 2-year steps for Brazil, SYS, and AIBL datasets, respectively). Second, significance testing and FDR between cell-type corrections were only applied for those cell-type groups that were significant in the main analysis.

We used a similar *Virtual Histology* analysis to assess the relationship between gene expression and cortical thinning in AD. For each ROI (n = 34), we modelled the effect of clinical diagnosis on cortical thickness with linear mixed models as implemented in *lme4*. The models also included magnetic field/site (ADNI/AIBL), age and sex as fixed effects and a subject identifier modelled as random intercepts. Significance testing was applied through permutation testing^24^, further corrected for multiple comparisons for the different cell-type panels (n = 9; FDR-corrected).

#### Independent Component Analysis

We used an ICA as implemented in FLICA^35,36^ with 70 components^80^ to derive modes of cortical thickness variability. For each IC, we initially tested the relationship with age using GAMM models. Following existing work^80,81^, only components with a strong *practical* significance (r^2^ > .15) were considered for further analysis. See **SI Methods** for more details.

#### Post-hoc analyses with the IC components

We tested the topological relationship between cortical thickness, T1w/T2w change during adolescence – as an index of myelination – and IC1 cortical weights using Pearson’s correlation. For T1w/T2w, we used a subset of LCBC participants (age = 8 – 40 years) with an available T2w sequence. We preprocessed the data using Human Connectome Project (HCP) processing pipeline^82^ (https://github.com/Washington-University/Pipelines) and summarized the T1w/T2w values – sampled at 70% from the WM/GM boundary^83^ - using the Desikan-Killiany parcellation. T1w/T2w values were fitted to age using splines and the mean derivative values between 15 and 21 were selected as the index of interest (age epoch where IC1 showed maximal decrements). See Grydeland and colleagues^84^ and **SI Methods** for more details. For IC2, scanner effects were tested using GAMM with scanner and sex as fixed effects, age as smoothing term and, participant as random effects. We used *geom_violin* and *geom_density* functions to visualize the relationship between scanner effect and scanner distribution through the lifespan with IC2 loading. For IC3, we performed a *virtual histology* analysis as described above, using IC3 cortical loadings as the phenotype of interest. Further, we used a co-expression analysis to test whether high-fidelity IC3 astrocytes, microglia and, CA1 pyramidal-specific genes were enriched for AD. Briefly, we created a co-expression data matrix based on available gene expression brain banks^26,84–87^ and overexpression for AD genes was then tested using an AD geneset curated by the DisGeNET database^89^. See **SI Methods** for more details.

## Acknowledgments

This work was supported by the Department of Psychology, University of Oslo (to K.B.W., A.M.F.), and the Norwegian Research Council (to K.B.W., A.M.F.). The project has received funding from the Coordenação de Aperfeiçoamento de Pessoal de Nível Superior - Brasil under Finance Code 001 (to A.P.J.), the European Research Council’s Starting Grant scheme under grant agreements 283634, 725025 (to A.M.F.) and 313440 (to K.B.W.) and the INTPART programme agreement 261577 (to K.B.W). Data used in the preparation of this article were partially obtained from the Australian Imaging Biomarkers and Lifestyle flagship study of ageing (AIBL) funded by the Commonwealth Scientific and Industrial Research Organisation (CSIRO), which was made available at the ADNI database (www.loni.usc.edu/ADNI). The AIBL researchers contributed data but did not participate in analysis or writing of this report. AIBL researchers are listed at www.aibl.csiro.au. Data collection and sharing for this project was funded by the ADNI (NIH Grant U01 AG024904) and DOD ADNI (Department of Defense award number W81XWH-12-2-0012). ADNI is funded by the National Institute on Aging, the National Institute of Biomedical Imaging and Bioengineering, and through generous contributions from the following: AbbVie, Alzheimer’s Association; Alzheimer’s Drug Discovery Foundation; Araclon Biotech; BioClinica, Inc.; Biogen; Bristol-Myers Squibb Company; CereSpir, Inc.; Cogstate Eisai Inc.; Elan Pharmaceuticals, Inc.; Eli Lilly and Company; EuroImmun; F. Hoffmann-La Roche Ltd and its affiliated company Genentech, Inc.; Fujirebio; GE Healthcare; IXICO Ltd.; Janssen Alzheimer Immunotherapy Research & Development, LLC.; Johnson & Johnson Pharmaceutical Research & Development LLC.; Lumosity; Lundbeck; Merck & Co., Inc.; Meso Scale Diagnostics, LLC.; NeuroRx Research; Neurotrack Technologies; Novartis Pharmaceuticals Corporation; Pfizer Inc.; Piramal Imaging; Servier; Takeda Pharmaceutical Company; and Transition Therapeutics. The Canadian Institutes of Health Research is providing funds to support ADNI clinical sites in Canada and the Saguenay Yout Study. Private sector contributions are facilitated by the Foundation for the National Institutes of Health (www.fnih.org). The grantee organization is the Northern California Institute for Research and Education, and the study is coordinated by the Alzheimer’s Therapeutic Research Institute at the University of Southern California. ADNI data are disseminated by the Laboratory for Neuro Imaging at the University of Southern California.

## Author Contributions

D.V.P, T.P., A.M.F, Z.P., and K.B.W. designed the study; A.P.J., G.S., and AIBL collected data; D.V.P, N.P., J.S., L.F., H.G., A.M.M., Y.P., and Ø.S. performed the analyses, created the figures. The paper was written by D.V.P, T.P. and, A.M.F with input from all the authors.

## Data availability

The authors will provide the code upon acceptance at https://github.com/LCBC-UiO/virtual_histology_2019.

