## Supplementary material for "Cellular correlates of cortical thinning throughout the lifespan": SI Methods

### Supplementary Information: Cellular correlates of cortical thinning throughout the lifespan

#### Supplementary Information: Methods

##### MRI sample. LCBC dataset.

The main sample was drawn from six Norwegian studies coordinated by the Center for Lifespan Changes in Brain and Cognition (LCBC; University of Oslo [UiO]): The Norwegian Mother and Child Cohort Neurocognitive Study[1]; Neurocognitive Development[2]; Cognition and Plasticity Through the Lifespan[3]; Constructive Memory[4]; Method of Loci[5]; and Neurocognitive plasticity[6]. The dataset was drawn on May 1, 2018. Next, we detail the main characteristics and the participants of each study.

*The Norwegian Mother and Child Cohort Neurocognitive Study.* This study consists of a subsample of the Mother-Child prospective study (MoBa)[7], by the Norwegian Institute of Public Health. Participants in the subsample underwent additional neuropsychological and neuroimaging assessments, coordinated by the center for LCBC. Parents enrolled in the Mother-Child study were contacted via mail for a request to join this subsample. Those who answered in written form were contacted and their children were added to the sample. Neuroimaging and Neuropsychological assessments took place in two Norwegian centers, in Oslo (Center for LCBC) and in Trondheim (St. Olav’s University Hospital, Norwegian University of Science and Technology [NTNU]). The project was approved by the Regional Committee for Medical and Health Research Ethics. Written informed consent was obtained from the parent/guardian for all participants and oral assent was given by participants at both time points. Participants were required to be fluent Norwegian speakers and have normal or corrected-to-normal vision and normal hearing. Exclusion criteria were history of injury or disease known to affect the central nervous system (CNS) function, including neurological or psychiatric illness, serious head trauma such as been unconscious, being under psychiatric treatment, use of psychoactive drugs known to affect CNS functioning, low birth weight (< 2500 g), magnetic resonance imaging (MRI) contraindications, and deemed free of brain injuries or pathological conditions. The study consists of two waves separated by 1.5 (SD = 0.2) years. Data acquisition took place between 2011 and 2014. MRI sequences were acquired with two 1.5 T Avanto scanners, located in Oslo and Trondheim. 720 scans were included in the sample from 466 unique individuals. Participants were aged between 4.1 and 12.1 (mean age = 7.4 [SD = 1.6]) years. See [1,8] for more details.

*Neurocognitive Development.* This study consists of a longitudinal sample of children and adolescents acquired at the center for LCBC. Typically developing children and adolescents aged between 8 and 19 years were recruited through newspaper ads and local schools. Written informed consent was obtained from all participants older than 12 years of age and from a parent of participants under 16 years of age. Oral informed consent was given by participants under 12 years of age. The study was approved by the Regional Ethical Committee of South Norway. Participants were required to be right-handed, fluent Norwegian speakers, have normal or corrected-to-normal vision and hearing, not have history of injury or disease known to affect CNS function, including neurological or psychiatric illness or serious head trauma, not be under psychiatric treatment, not use psychoactive drugs known to affect CNS functioning, not have had complicated or premature birth, not have MRI contraindications, and deemed free of brain injuries or pathological conditions. The study consists of three waves. Participants were re-scanned approximately after 3.5 years. Data acquisition took place between 2007 and 2016. MRI sequences were acquired either with a 1.5 T Avanto or a 3.0 T Skyra scannerp located at the Oslo University Hospital. 665 valid scans were included in the sample from 326 unique individuals. Participants' age ranged between 8.1 and 26.7 years (mean age = 17.4 [SD = 4.3]). See [2,9] for more details.

*Cognition and Plasticity through the Lifespan.* The study consists of a large longitudinal study where cognitively healthy adults spanning the whole age span underwent MRI scanning and neuropsychological evaluation. The sample was collected at the center for LCBC. Volunteers were initially recruited by newspaper advertisements and later contacted by mail for follow-ups. The study was approved by the Regional Ethical Committee of South Norway, and written informed consent was obtained from all participants prior to the examinations. Exclusion criteria included history of injury or disease known to affect the CNS function, including neurological or psychiatric illness or serious head trauma, being under psychiatric treatment, use of psychoactive drugs known to affect CNS functioning, and MRI contraindications. Participants were deemed free of dementia, moderate depression (≥25 on the Mini-Mental State Examination (MMSE)[10], Beck Depression Inventory (BDI; ≤21)[11] and significant brain injuries or conditions. Additionally, participants were required to be right-handed, native Norwegian speakers, have normal or corrected-to-normal vision, and score ≥85 on the Wechsler Abbreviated Scale of Intelligence (WASI)[12]. The study consists of four waves. New participants were recruited in waves 1 and 3. Participants were scanned up to 11.1 years after the initial scan. The interval between waves was approximately 3.5, 4.4 and, 1.7 years, respectively. Data acquisition took place between 2006 and 2018 (ongoing). MRI sequences were acquired with a 1.5 T Avanto and with 3.0 T Skyra and Prisma scanners located at the Oslo University Hospital. 1284 valid scans were included in the sample from 732 unique individuals. Participants' age at test ranged between 20.0 and 89.4 years (mean age = 48.4 (SD = 18.0)). For more details see [3,13,14].

*Constructive Memory.* The study consists of a cross-sectional study where cognitively healthy adults spanning the whole age span underwent an fMRI source-item memory task. The protocol also included MRI scanning and neuropsychological evaluation. The sample was collected at the center for LCBC. The project is nested within the *Cognition and Plasticity through the Lifespan* project. All participants gave written informed consent and the study was approved by the Regional Ethical Committee of South Norway. Participants were screened through health and neuropsychological interviews. Exclusion criteria included neurologic or psychiatric disorders, chronic illness, premature birth, learning disabilities, handedness or, current use of medicines known to affect nervous system functioning. Participants were also excluded based on neuropsychological evaluation criteria: score <26 on the MMSE, score of ≥16 in the BDI, score <85 on the WASI, and a T-score of ≤30 on the California Verbal Learning Test II—Alternative Version (CVLT II)[15] immediate delay and long delay. Participants were required deemed free of significant brain injuries or conditions. The study consists of a single wave. Data acquisition took place between 2013 and 2015. MRI sequences were acquired with a 3.0 T Skyra scanner located at the Oslo University Hospital. 165 participants were included in the sample. Participants were aged between (mean age = 35.0 [SD = 15.2] years; age range = 19.4 – 76.8) years. See [4,16] for more details.

*Method of Loci.* This study consists of an eight-week memory training experiment focused on improving verbal recall memory by implementing the mnemonic technique method of loci. Participants were randomly assigned to two groups, either the intervention group or a control group serving as passive controls. Participants were scanned three times: pre and post-training and a follow-up after 5 years. Volunteers were recruited through a local newspaper ad and screened by a structured interview before inclusion. Cognitive assessments and the memory training program were conducted at the center for

LCBC. All participants gave informed consent, and the study was approved by the Regional Ethical Committee of South Norway. All included participants reported to be right-handed, native Norwegian speakers, not concerned about their own memory function, not using medications known to interfere with cognitive function and having no diseases known to affect the CNS including severe hypertension or diabetes. Exclusion criteria included: MMSE < 26, WASI < 85, and Geriatric Depression Scale (GDS) > 11[17]. Further, participants with total learning and/or recall scores on the CVLT II of 2 SD or more below population norm were also excluded. Furthermore, all MRI scans were subjected to a radiological evaluation by a neuroradiologist and were required to be deemed free of significant injuries or conditions. Data acquisition for waves 1 and 2 took place between 2007 and 2008 while the third wave was acquired in 2013. MRI sequences were acquired with a 1.5 T Avanto scanner located at the Oslo University Hospital. 174 observations from 70 unique participants are included in the sample. Participants age at the first time point was 62.3 (SD = 9.5; age range 41.9 – 79.1 years) years. Participants were re-scanned ≈ after 0.2 and 5.4 years at wave 2 and 3, respectively. See [5,18] for more details.

*Neurocognitive plasticity*. The study consists of an experimental project of memory training with the method of loci. The study includes two groups of participants (young and old) that underwent an ABAB design where a batch in each group started with a resting condition, and the other started with memory training (after a baseline test). The study also includes an active group without memory training. The participants were scanned 6 times, five of them before/after a block of training while the sixth time point consisted of a follow-up ≈ 2 years after the intervention. All procedures were approved by the regional ethical committee of Southern Norway, and written consent was obtained from all participants. Participants were recruited through newspaper and web page adverts and were screened with a health interview. Participants were required to be either young or older (in or around their 20s or 70s, respectively) healthy adults, right-handed, fluent Norwegian speakers, and have normal or corrected to normal vision and hearing. Exclusion criteria included history of injury or disease known to affect CNS function, including neurological or psychiatric illness or serious head trauma, being under psychiatric treatment, use of psychoactive drugs known to affect CNS functioning, and MRI contraindications. Moreover, for inclusion in the present study, participants were required to score ≥ 26 on the MMSE and have scores within the normal range (≥2 SD below mean) for age and sex on the 5-min delayed recall subtest of the CVLT II. All participants further had to achieve a WASI score > 85. Participant scans were evaluated by a neuroradiologist and deemed free of significant injuries or conditions. Data acquisition took place between 2013 and 2018 (ongoing). MRI sequences were acquired with a 3.0 T Skyra scanner located at the Oslo University Hospital. 956 observations from 229 unique participants are included in the sample. Mean age for the young and old group of participants was 27.0 (SD = 3.3; age range = 20.1 – 35.0) and 74.0 (SD = 3.0; age range = 68.7 – 84.0) years. Participants were re-scanned ≈ after 0.20, 0.41, 0.63, 0.84 and 2.9 years after the initial scan, respectively. See [19,20] for more details.

##### MRI sample. Replication and AD datasets.

Three additional datasets were used to replicate the main findings of the lifespan virtual histology analysis. Each replication dataset consists of cognitively healthy participants spanning the age ranges where significant expression – thinning correlations were found. See **Figure S2** and **Table S5** for graphical representation and descriptive information on each cohort.

*Brazilian High-Risk Cohort.* The replication for the older adults was carried out with the Brazilian High-Risk Cohort dataset, designed to better understand the development trajectories of psychopathology and mental disorders in children and adolescents. It consists of a large longitudinal MRI sample of children and adolescents that took place in two Brazilian centers (Porto Alegre and São Paulo)[21]. The enrollment for the screening phase was conducted at public schools during the early registry days. The final sample was enriched for individuals with high risk for developmental psychiatric disorders. This study was approved by the ethics committee of the University of São Paulo. Written consent was obtained from parents of the participants as well as from those participants that were able to read, write and understand the written consent, else a verbal agreement was obtained. For a subsample of the cohort, MRI data was acquired. See [21] for details.

For the replication sample, we selected those observations with MRI structural information. The cohort consisted of 809, unique participants (350 females), acquired in two different waves (n = 724, 450). The mean interscan interval was 3.8 (SD = 0.38) years. The mean age of the participants was 12.0 (SD = 2.6; age range = 6.9 – 18.9) years. The MRI dataset was acquired in two centers (Porto Alegre, São Paulo) equipped with 1.5 T scanners.

*Saguenay Youth Study - Parents cohort (SYS).* The replication for the middle-aged adults was carried out with the parent arm of the Saguenay Youth Study (SYS). The SYS goal was to understand trans-generational models of developmental cascades and its relationship with the emergence of common chronic disorders[22]. The full cohort consists of a longitudinal community-based sample of adolescents and their parents from the Saguenay Lac Saint Jean region (Quebec, Canada). Adolescents were recruited in local high schools and the main exclusion criteria were MRI contraindications and serious conditions likely to affect the brain or heart development. All participants included in the study are of single ethnicity, i.e. White Canadians of French descent. The study was approved by the Chicoutimi Hospital Research Ethics Committee. Written informed consent and assent were obtained from the parents and adolescents, respectively. See [22,23] for further details.

For the replication sample, we considered MRI data from the parent arm of the SYS cohort. Thus, the replication dataset consisted of 548 unique participants (300 females) with a single scan available per participants. The mean age of the participants was 49.4 (SD = 5.0; age range = 36.4 – 65.4) years. The MRI dataset was acquired in a single 1.5 T scanner.

*Australian Imaging, Biomarker & Lifestyle Flagship Study of Ageing (AIBL).* The replication for the older adults was carried out with the AIBL study. The AIBL study took place in Australia (Perth and Melbourne), beginning in 2006, with the aim to study which biomarkers, cognitive characteristics, and health and lifestyle factors determine the development of Alzheimer's disease (AD). It consists of a 4.5 years longitudinal prospective study with neuroimaging and neuropsychological assessments every 1.5 years. The cohort includes a sample of >1,100 participants aged > 60 years. At baseline, participants were either healthy individuals, individuals categorized as mild cognitive impairment (MCI) or AD patients. Cognitively healthy participants also included volunteers that expressed subjective concern about their memory function. See [24,25] for further details.

Exclusion criteria in the individuals that volunteered included: non-English speaking, non-AD neurologically relevant history (e.g., traumatic brain injury), schizophrenia; depression (GDS > 5);

Parkinson's disease; cancer within the last 2 years; symptomatic stroke; uncontrolled diabetes; or current regular alcohol use exceeding standard drinks per day. The study was approved by and complied with the regulations of the institutional research and ethics committees of Austin Health, St. Vincent's Health, Hollywood Private Hospital, and Edith Cowan University[24]. All participants provided written informed consent prior to participating in the study. All participants enrolled in AIBL underwent a thorough medical and neuropsychological evaluation at each assessment and this information was used to classify the clinical disease stage. A clinical review panel reviewed all available medical, psychiatric and neuropsychological information to confirm the cognitive health of individuals enrolled in the cognitively normal group. In this clinical review panel, participants' cognitive and functional status was rated using the Clinical Dementia Rating scale (CDR)[26]. For this study, individuals classified as CDR = 0.5 were considered to have very mild dementia or MCI, while those with CDR = 0 were classified as cognitively normal.

For the replication sample, we only considered cognitively normal participants. Participants that converted to MCI or AD at later visits were still considered. Data for participants was thus considered only when CDR = 0 and MMSE > 25. Drop-out and conversion were not further modeled. Thus, the AIBL replication dataset consisted of 739 unique observations, from 435 unique participants (246 females) acquired in four waves (n = 417, 131, 104 and, 87, respectively). Each wave was acquired every 15 months (spanning up to 4.5 years). The mean age of the participants was 72.6 (SD = 6.5; age range = 60.0 – 92.0) years. The MRI images were acquired in three different 1.5 T scanners.

For the AD dataset we considered all the available MRI observations with clinical diagnosis status of cognitively healthy (HC), Mild Cognitive Impairment (MCI) or Alzheimer's Dementia (AD). The AIBL AD dataset (for replication of Virtual Histology throughout the AD continuum) consisted of 1069 MRI observations with 631 unique participants (281 females; age = 73.6 [6.8] years; age range = 55.0 – 95.7). Of these observations, 785 corresponded to HC individuals, 149 to MCI patients, and 135 to AD patients.

*Alzheimer's Disease Neuroimaging Initiative (ADNI).* Cortical thinning patterns across the AD continuum were characterized by using the ADNI database (discovery sample; adni.loni.usc.edu). The ADNI was launched in 2003 as a public-private partnership, led by Principal Investigator Michael W. Weiner, MD. The primary goal of ADNI has been to test whether serial MRI, positron emission tomography (PET), other biological markers, and clinical and neuropsychological assessment can be combined to measure the progression of MCI and early AD. For up-to-date information, see www.adni-info.org. ADNI is a multisite longitudinal study, with 63 (57 recruiting) sites across the USA and Canada. ADNI enrolls participants between the ages of 55 and 90. It currently consists of 4 phases (ADNI, ADNIGO, ANDI2, ADNI3)[27,28] of which the later, ADNI3, is currently ongoing. For our dataset, we considered ADNI1, ADNIGO and ADNI2 scans. Each wave consists of a 4 years longitudinal prospective study which – amongst others – include neuroimaging assessments every 12 months. The original ADNI dataset followed-up 200 HC, 400 MCI and, 200 mild AD participants. ADNIGO included normal controls and MCI participants from ADNI1 and 200 new MCI participants. ADNI2 included CN and MCI rollover participants in addition to new 650 new participants (150 HC, 300 MCI; 200 mild AD)

Basic inclusion and exclusion criteria are thoroughly described elsewhere[29]. It includes an age range between 55 and 90 years of age and a study partner able to provide an independent evaluation of functioning. Subjects could speak either English or Spanish. All subjects had to be willing to undergo all test procedures including neuroimaging and longitudinal follow-up. At least 20% of the subjects at each site had to be willing to undergo 2 lumbar punctures spaced 1 year apart. Psychoactive medications that were believed to possibly affect cognitive function were excluded. No evidence of serious ischemic stroke (Hachinski Ischemic Score ≤ 4), stable medications for 4 weeks prior to screening and, GDS < 6 were also required. In addition, individuals required visual and auditory acuity enough for neuropsychological testing; good general health, no medical contraindications to MRI and 6 grades of education/work history. Subjects needed to sign an informed consent. The protocols were approved by the corresponding ethical committees.

For the ADNI dataset, we considered all the available MRI observations with available clinical diagnosis status of HC, MCI or AD. The ADNI dataset (for Virtual Histology analysis in AD) consisted of 6680 MRI observations with 1725 unique participants (774 females; age = 75.1 [7.2] years; age range = 54.4 – 95.6). Of these observations, 2075 corresponded to HC individuals, 3097 to MCI patients, and 1508 to AD patients.

#### MRI acquisition parameters for the LCBC dataset.

For *Avanto1* and *Avanto2* (Siemens Medical Solutions, 1.5 T), data were acquired using a 12-channel head coil. Two identical 3D T1-weighted magnetization prepared rapid gradient echo (MPRAGE) sequences were obtained with the following parameters: TR/TE/TI = 2400 ms/3.61 ms/1000 ms, FA = 8°, acquisition matrix = 192 × 192, FOV = 192, 160 sagittal slices with voxel size = 1.25 × 1.25 × 1.2 mm. Skyra (Siemens Medical Solutions, 3.0 T) data were collected using a 24-channel coil. Anatomical T1-weighted MPRAGEs consisted of 176 sagittally oriented slices obtained using a turbo field echo pulse sequence (TR/TE/TI = 2300 ms/2.98 ms/ 850 ms, FA = 8$^{o}$, acquisition matrix = 256 × 256 with voxel size = 1×1×1 mm). Prisma (Siemens Medical Solutions, 3.0 T) data were collected using a 32-channel coil. Anatomical T1-weighted MPRAGEs consisted of 208 sagittally oriented slices obtained with the following parameters: TR/TE/TI = 2400 ms/2.22 ms/ 1000 ms, FA = 8$^{o}$, acquisition matrix = 256 × 256 with voxel size = 0.8 × 0.8 × 0.8 mm, GRAPPA = 2. For most children < 9 years, integrated parallel acquisition techniques (iPAT) were used, acquiring multiple T1 scans within a short scan time, to discard scans with residual movement and to average the scans with sufficient quality.

#### MRI acquisition parameters for the replication samples and AD datasets.

*Brazilian High-Risk Cohort.* Imaging data were acquired at the University of São Paulo's Institute of Radiology and the Santa Casa of Porto Alegre using a 1.5 T MRI Signa HDx and a GE Sigma HD scanners (General Electric), respectively. The T1-weighted acquisition parameters were as follows: TR = 10.916 ms, TE = 4.2 ms, thickness = 1.2 mm, 156 slices, flip angle = 15°, NEX = 1, matrix size = 256 × 192, FOV = 245 mm, and bandwidth = 122.109.

*Saguenay Youth Study - Parents cohort (SYS).* Imaging data were acquired at the MR Clinic in Chicoutimi using a 1.5 T Avanto (Siemens Medical Solutions). Three-dimensional (3D) radio frequency (RF)-spoiled gradient-echo scan with 140-160 slices was acquired with TR = 25ms, TE = 5ms, flip angle = 30°, voxel size = 1.0 x 1.0 x 1.0.

*Australian Imaging, Biomarker & Lifestyle Flagship Study of Ageing (AIBL).* The MRI dataset was acquired in a 1.5 T (Avanto) and two 3.0 T scanners (Verio, TrioTim) (all Siemens Medical Solutions) located at Perth and Melbourne. Sagittal T1-weighted MRI scans were acquired using a standard 3-dimensional magnetization-prepared rapid gradient echo (MPRAGE) sequence. The T1-weighted parameters were as follows: 160 slices, voxel size = 1.0 x 1.0 x 1.2, matrix size = 240 x 256, TR = 2300ms, TE = 2.98 ms, flip angle = 9°, TI = 900ms.

*Alzheimer's Dementia Imaging Initiative (ADNI).* Most of the ADNI1 and ADNIGO-rollover participants were scanned using 1.5T scanners while ADNI2 participants were exclusively scanned with 3T equipment. Sagittal T1-weighted MRI scans were acquired using standard 3-dimensional MPRAGE sequences. The T1-weighted parameters slightly varied due to differences in scanners and vendors (Philips, GE, Siemens) and evolution throughout the different ADNI waves. See more information at http://adni.loni.usc.edu/methods/mri-tool/mri-analysis/.

#### MRI longitudinal FreeSurfer v.6.0. stream

First, data entered the cross-sectional pipeline, which includes removal of non-brain tissue, Talairach transformation, intensity correction, tissue and volumetric segmentation, cortical surface reconstruction and cortical parcellation. The pipeline yields a reconstructed surface map for each person at each point. To extract more reliable thickness estimates, the reconstructed images were fed to the longitudinal stream. Specifically, an unbiased within-subject template space and image are created using robust, inverse consistent registration [30]. Several processing steps, such as skull stripping, Talairach transforms, atlas registration, as well as spherical surface maps and parcellations, are then initialized with common information from the within-subject template, significantly increasing the reliability and the statistical power of the cortical thickness estimates.

#### Modifications of the MRI processing pipeline in the replication and the AD datasets.

The replication datasets were all processed using the same FreeSurfer pipeline as described in the main text with the exception of minor variations, listed next. 1) For the replication datasets, Freesurfer version 5.3 was employed for cortical reconstruction of the images. 2) The *Brazilian High-Risk* and the SYS MRI datasets were processed at a processing core located in Toronto, respectively. 3) The SYS and the *Brazilian High-Risk* datasets were fed into the cross-sectional pipeline, instead of the longitudinal pipeline. ADNI (discovery) and AIBL (replication) AD datasets, were also processed using the same FreeSurfer pipeline as described in the main text. The only variation was that datasets were not fed to the longitudinal pipeline to reduce inter-diagnostic biases in each individual's FreeSurfer template.

#### Consistency of inter-regional profiles in gene expression.

Only consistent genes were retained for Virtual Histology analysis. Gene consistency was assessed through two different steps. In stage 1, we calculated the mean Spearman correlation between each of the six donor's profiles and the median (group) profile for each gene. Using this metric, the median profile provides a good approximation across the donors for 39.7% of the assayed genes (rho > 0.446 corresponding to one-sided $p < 0.05$ derived from random simulations of donor expression profiles)[31]. In stage 2, we relied on the BrainSpan Atlas, which provides gene expression data in the developing human brain (www.brainspan.org)[32] as previously implemented[33]. We limited the samples to age > 12 years (n = 9 donors) and downloaded gene expression values obtained in 11 cortical regions included in the BrainSpan atlas that are homologous to those on the Desikan-Killiany parcellation employed in the Allen Human Brain Atlas. Next, we compared the similarity between the regional profile between the Allen Human Brain Atlas and the BrainSpan Atlas across the 11 cortical regions available in both atlases. Only genes that showed a correlation between the two profiles higher than r = 0.52 (one-sided test $P < 0.05$) were retained.

#### Virtual Histology. Cell type – cortical thinning relationship for cell type subclasses.

Panels of Subclass Cell-Specific Marker Genes. In addition to the main nine classes of cells described above, we explored the association between cortical thinning and expression for specific subclasses of cells as obtained from Zeisel et al.[34]. Following previous work[35], we log-transformed the RPKM values, added 1 and, Z-standardized the values across cells. Genes with average standardized expression levels > 2 SD in a given transcriptomic cell type subclass were considered enriched. Cell subclass panels of genes were obtained by filtering the n = 2,511 genes and restricting each subclass panel up to 21 marker genes.

Correlation with cell type subclasses. For each developmental period of interest – namely, childhood, middle-age and older age - the cortical profiles of thinning were averaged to a single phenotype per period. Then, we assessed the correlation between thinning profiles in childhood, middle-aged and old adulthood and gene expression profiles for each subtype of cell as described before – excluding the multiple comparisons correction for age. In addition, the different subclasses of interest were ranked - i.e. from higher to lower expression - thinning correlation - to better evaluate whether different cell type subclasses drive the thinning-expression relationship in the childhood and the aging periods.

#### Linked ICA – Modes of cortical thickness variation analysis

To obtain modes of cortical thickness variation throughout the lifespan we used a linked Independent Component Analysis (ICA) as implemented in FLICA (http://fsl.fmrib.ox.ac.uk/fsl/fslwiki/FLICA)[36,37]. The linked-ICA approach provides a data-driven decomposition of the images into spatial components characterizing intersubject variability. ICA approaches are able to model data into a set of – maximally independent; not orthogonal - interpretable features, some of them linked to biophysically plausible underlying mechanisms, which can additionally be linked to external variables such as age. Here, each spatial component represents a mode of variation of cortical thickness variation across the n = 4.004 observations. Note that the analysis does not permit the introduction of random effects (i.e. to model repeated observations per participant). However, if repeated observations are removed, we obtained similar modes of cortical thickness variations across the lifespan. We ran the linked ICA decomposition – as implemented in FLICA[36,37] - with 70 components[38]. For each independent component, we initially tested the relationship with age using generalized additive mixed models (GAMM). Only components with practical significance with age ($r^2 > .15$ as defined by Douaud and colleagues[38]) were considered for further analysis. Three components (IC1, IC2 and, IC3) achieved practical age significance and were inspected further.

**Post-hoc analysis with selected IC components**

Relationship between IC1 cortical weights and T1w/T2w changes during adolescence

The T1w/T2-weighted (T2w) preprocessing and analysis of the age trajectories has been described in detail in Grydeland and colleagues[39]. See below for a summarized description of the methods.

*Participants.* The T1w/T2w dataset included neuroimaging data for 263 participants (136 females; mean age = 19.4 [7.9] years; age range = 8.2 - 40.0). The sample was drawn from the first wave of the Neurocognitive Development[2] and the Cognition and Plasticity Through the Lifespan[3] LCBC's datasets. See characteristics, ethics approval and, exclusion criteria above (section: *MRI sample. LCBC dataset*). The participants corresponded to a subsample of participants and observations used in the main LCBC sample. An available T2w scan after quality control and age < 40 years were additional inclusion criteria for this sample.

*MRI Acquisition and preprocessing.* Scans were acquired in the 1.5T Siemens Avanto scanner located at Oslo University Hospital. For each participant, the T1w and the T2w scans were acquired in the same session. The T1w MPRAGE sequence parameters were as described above (TR/TE/TI = 2400 ms/3.61 ms/1000 ms, FA = 8°, acquisition matrix = 192 × 192, FOV = 192, 160 sagittal slices with voxel size = 1.25 × 1.25 × 1.2 mm). The T2w volumes were acquired using a 3D T2w sampling perfection with applicationoptimized contrasts using different flip angle evolutions (SPACE) sequence with the following parameters: TR/TE 3390 ms/388 ms, variable FA, FOV = 256 mm, 1 mm isotropic voxels. Participants were scanned either with a 204 x 256 x 176 matrix or with a 256 x 256 x 176 matrix. Differences in T1w/T2w due to matrices size were estimated via robust regression and removed from T1w/T2w estimates as described in Grydeland and colleagues[39].

For each participant, T1w/T2w ratio maps were created using the Human Connectome Project (HCP) processing pipeline[40] (https://github.com/Washington-University/Pipelines), which included T1w processing with the cross-sectional Freesurfer 5.3 suite (described above). The present pipeline deviates from the HCP pipeline in that we did not perform gradient distortion nor readout distortion correction. The T2w image was registered to the T1w image by using Freesurfer's *bbregister*, a within-subject, cross-modal registration using a *boundary-based cost* (BBC) function constrained to be 6 degrees of freedom (rigid body)[44]. The resulting linear transform was applied by using FSL's *applywarp* tool using spline interpolation in order to minimize the white matter and cerebrospinal fluid contamination of GM voxels[45]. The T1w volume was divided on the aligned T2w volume, creating a T1w/T2w ratio volume. To estimate regional T1w/T2w, we used the Desikan-Killiany atlas to divide each cerebral hemisphere into 34 regions. We then sampled T1w/T2w values at 70% from the WM/GM boundary[46].

*T1w/T2w signal change in adolescence.* Age trajectories were fitted with penalized splines[47,48]. Eight piecewise cubic B-spline basis functions were used to fit a smooth, non-linear curve from the weighted sum of these functions. For each region, we calculated the derivatives of the T1w/T2w age curve using an approximation by finite differences. Finally, we took the mean derivative between 15 and 21 years of age – the age epoch where the IC1 component has the steepest curve – as the measure of interest and used it as a putative measure of myelination during development. We performed a Pearson's correlation (n = 34 Desikan-Killiany ROIs) to assess the relationship between IC1 cortical weights and T1w/T2w changes during development.

*<u>Co-expression analysis between high-fidelity IC3 cell-specific genes and Alzheimer's Disease genes.</u>*

*Co-expression matrix.* To study the co-expression of high-fidelity IC3 cell-specific genes we followed the co-expression pipeline described by Sliz and colleagues[49]. Briefly, we created a co-expression matrix based on cortical brain samples from n = 572 unique brain donors acquired from five human (Allen Human Brain Atlas[50,51], BrainCloud[52,53], Braineac[54], BrainSpan[55], and GTEx[56]) between the ages of 0 and 102 years of age at death[57]. Gene expression was quantified using microarrays for the AHBA, BrainCloud, and BrainEAC while RNA sequencing was used for BrainSpan and GTEx. For each database and gene, Entrez gene symbols were updated while values were log10 transformed and scaled within each sampled region. After this pre-processing stage, the gene expression was combined across all databases. Only genes with expression values from all five databases were included in the final curated expression database (16245 genes). For each of the 16245 genes, we tested co-expression using linear mixed models (adjusted for age, sex, and repeated measures as random effect). For each of the tested genes, co-expression genes were determined by the top 1% as determined by smallest p-values of the gene effects term.

Enrichment analysis for *high-fidelity IC3 cell-specific co-expressed genes.* Gene co-expression analysis was carried for the high-fidelity IC3 cell-specific genes. The gene sets were obtained by 1) selecting those cell type panels whose expression was associated with lifespan cortical thinning (IC3; i.e. microglia, astrocyte and CA1 pyramidal-specific genes) and 2) filtering those genes that were significantly related to the cortical weights for IC3 ($p < .05$, n = 34 ROIs, FDR corrected). Co-expression genes were obtained for the following high-fidelity IC3 gene sets (microglia n = 24; astrocyte n = 26 and, CA1 pyramidal n = 36). The resulting set of co-expressed genes (microglia n = 161; astrocyte n = 178 and, CA1 pyramidal n = 292) were used in a gene ontology (GO) enrichment analysis for AD as curated in the DisGeNET database[58]. Significance was controlled FDR for multiple comparisons (n = 3 co-expressed sets of genes).

#### Supplementary Information: Results

##### Expression – thinning correlates (Virtual Histology). Effects of different fitting parameters.

We repeated the virtual histology approach (i.e. cell-specific gene expression – cortical thinning correlation) from thinning estimates with varying fitting parameters. Specifically, we modified the number of knots, at 6, 7, 8, 9, 10, 12, 15, 20, 30, and 40 knots. The knots are locations where polynomial curves are joined. The fewer the knots, the smoother the spline curve is and vice versa. A high number of knots likely leads to overestimated trajectories of cortical thickness and thinning, by means of the $1^{st}$ derivative. See results in **Figure S1**. The mirrored pattern at both ends of the lifespan remained evident regardless of the fitting parameters, though microglia gene expression tends to show greater negative correlations with thinning at the very old age. The association of astrocyte-specific gene expression with thinning profiles at middle-age was evident at low fitting complexity and at very high fitting complexity but disappeared when fitted at 12 and at 15 knots.

##### Expression – thinning correlates (Virtual Histology). Replication samples

*Brazilian High-Risk Cohort.* We used the Brazilian High-Risk Cohort to replicate the virtual histology results in the youngest segment of the lifespan (age range of the dataset = 7 to 19 years). As in the original sample, we found an association between astrocytes, microglia, and CA1 pyramidal-specific gene expression and the cortical thinning profiles in childhood and adolescence (i.e. more expression, less thinning). The expression – thinning correlation coefficients for astrocytes and CA1 pyramidal cells were maximal at younger ages (r = .17 and .12 respectively) decreasing to non-significant values hereafter (CA1 pyramidal cells showed a significant association again at age 17). Microglia-specific gene expression showed a more stable relationship with thinning throughout development (correlation coefficient range = .13 to .15), reaching significance at age 9, 13 and, 15 years of age. The expression – thinning correlation for S1 pyramidal and oligodendrocytes tended to exhibit a negative pattern at later adolescence (> 15 years) reaching a negative relationship of r ≈ .15 (i.e. more gene expression, steeper thinning).

*SYS.* We used the parental arm of the SYS cohort to replicate the virtual histology results in the middle-aged segment of the lifespan (age range of the dataset = 34 to 65 years). As in the original sample, we found an association between astrocyte-specific gene expression and the cortical thinning profiles at middle-age (i.e. more expression, reduced thinning). The correlation coefficients were maximal between 34 and 47 years of age (r = .19) and decreased hereafter reaching non-significant levels (r ≤ .1) at age 60. The expression-thinning correlations for all the remaining cell types were lower than those observed for astrocytes.

*AIBL.* We used the AIBL dataset to replicate the virtual histology results in the older segment of the lifespan (age range of the dataset = 60 to 90 years). As in the original sample, we found a significant association between we found an association between astrocytes, microglia, and CA1 pyramidal-specific gene expression and the cortical thinning profiles in older age (i.e. more expression, steeper thinning). The expression-thinning correlation coefficients are relatively stable across older age, between -.14 and -.26

peaking around 70 years of age. The expression – thinning correlation for the remaining cell types is lower than those observed for astrocytes, microglia and, CA1-pyramidal cell types (between r = -.13 - .1).

#### Virtual Histology. Cell type – cortical thinning relationship for cell type subclasses.

Further, we tested whether different subclasses of astrocytes, microglia, and CA1 pyramidal cells could account for the cortical thinning – gene expression relationship in the childhood and the aging periods. We restricted the analyses to three developmental periods corresponding to the inter-regional patterns of cortical thinning in childhood (5 – 9 years), middle-age (35 – 41 years) and older age (65 – 67 years). The results showed that the same cell type subclasses are associated with thinning profiles during childhood and older adulthood (but – as expected based on the original analysis - with the opposite expression - thinning direction) (**Table S3**). For CA1 pyramidal cells, only expression for the *CA1Pyr2* subclass was associated with young and old cortical thinning profiles.

#### Supporting information

See https://athanasiamo.shinyapps.io/Virtual_histology_2019/ for supporting information and interactive visualization of the results.

#### Supplementary Figures

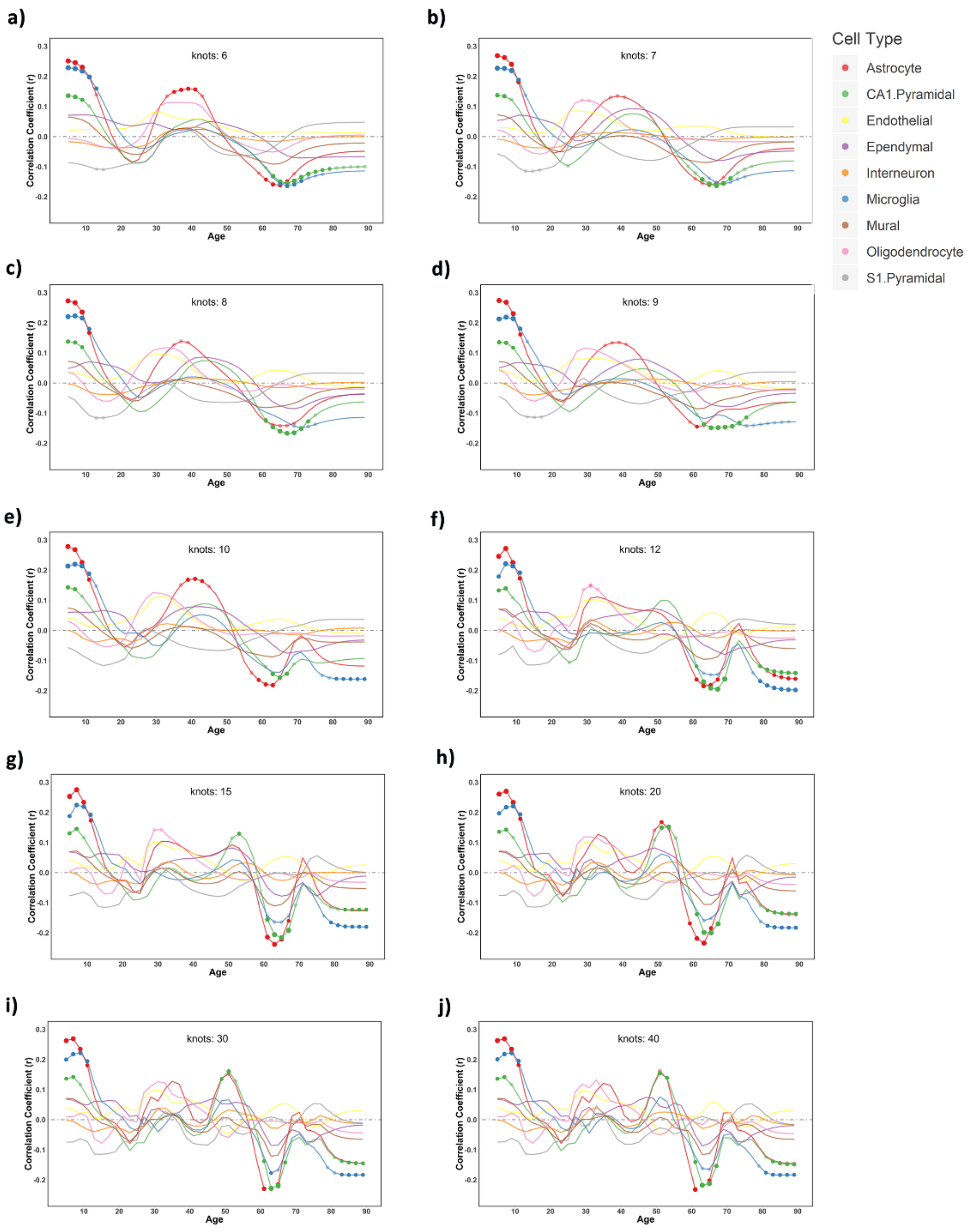


***Figure S1.*** *Virtual Histology across the lifespan – cortical thickness trajectories fitted with different parameters. Correlation coefficients between the cortical thinning profile across the lifespan and mean regional profiles of gene expression levels in each of the 9 cell types. For each panel, cortical thickness trajectories have been fitted with different levels of complexity. In each plot, the x-axis indicates age while*

*the y-axis indicates the correlation of thinning with cell type-specific gene expression as derived from the Allen Human Brain Atlas. Values above 0 represent a relationship of gene expression profiles with reduced thinning - or thickening - while values below 0 represent a relationship with steeper cortical thinning. Circles indicate a significant relationship ($p < 0.05$, permutation inference $n = 10000$ iterations) after Bonferroni correction for multiple comparisons along the lifespan (semi-transparent circles) and after additionally applying FDR-adjustment for testing multiple cell types (opaque circles). The size of the circle represents significance. Thinning profiles were derived from the LCBC dataset. ($n = 4004$ observations).*

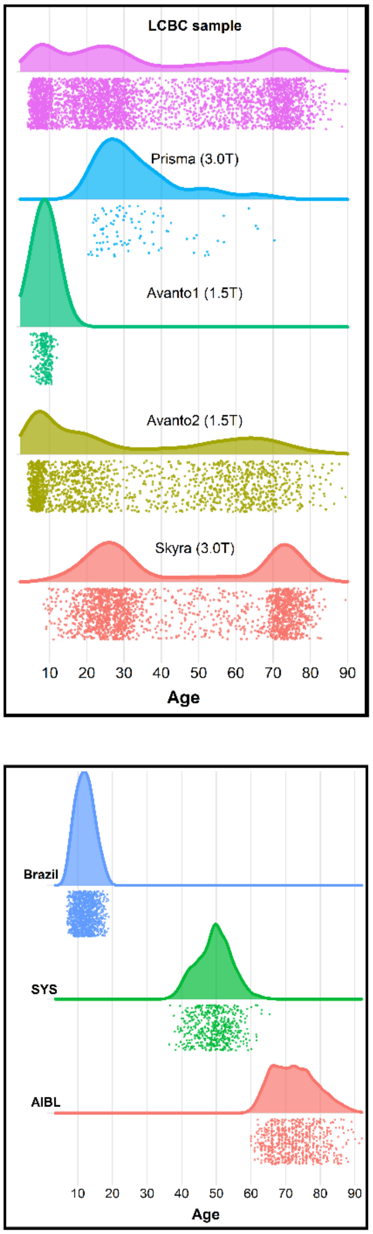


***Figure S2.*** *Distribution of observations across the lifespan. The upper panel shows the distribution of MRI observations for the LCBC dataset. The distribution of MRI observations is grouped by scanner. The lower panel shows the distribution of MRI observations for the three replication datasets.*

#### Supplementary Tables

| Site | Observations | Participants | Obs./participant (mean) | Sex (female:male) | Age (SD) | Age range |
|---|---|---|---|---|---|---|
| **LCBC (all)** | 4,004 | 1,899 | 2.1 | 1,135:764 | 37.4 (25.3) | 4.1 - 89.4 |
| **Skyra 3.0 T** | 1,934 | 1,081 | 1.8 | 684:397 | 46.0 (23.3) | 8.5 - 89.4 |
| **Avanto1 1.5 T** | 240 | 135 | 1.8 | 67:69 | 8.6 (1.4) | 4.9 - 12.1 |
| **Avanto2 1.5 T** | 1,737 | 875 | 2.0 | 470:405 | 32.1 (25.1) | 4.1 - 89.4 |
| **Prisma 3.0 T** | 93 | 93 | 1 | 68:25 | 33.5 (11.3) | 20.5 - 70.3 |

***Table S1***. *MRI sample descriptives. Main sociodemographic information of the MRI sample, grouped also by scanner. Obs. = Observations.*

| | | Young | | | Middle-aged | | | Old | | |
|---|---|---|---|---|---|---|---|---|---|---|
| **Cell type** | **Genes(n)** | **r** | **p** | **p (fdr)** | **r** | **p** | **p (fdr)** | **R** | **p** | **p (fdr)** |
| **Astrocyte** | 54 | .19 | <.001 | <.001 | .15 | <.001 | .002 | -.11 | .005 | .02 |
| **CA1.Pyramidal** | 103 | .10 | .001 | .004 | .06 | .42 | .95 | -.14 | <.001 | .001 |
| **Endothelial** | 57 | .02 | .58 | .66 | .06 | .35 | .95 | .02 | .72 | .81 |
| **Ependymal** | 84 | .07 | .04 | .08 | .04 | .79 | 1 | -.07 | .05 | .10 |
| **Interneuron** | 100 | -.03 | .46 | .64 | .03 | .92 | 1 | -.01 | .86 | .86 |
| **Microglia** | 48 | .18 | <001 | <.001 | .02 | .90 | 1 | -.15 | <.001 | .002 |
| **Mural** | 25 | .04 | .50 | .64 | .01 | .91 | 1 | -.06 | .32 | .57 |
| **Oligodendrocyte** | 60 | -.02 | .66 | .66 | .11 | .005 | .02 | -.02 | .66 | .81 |
| **S1.Pyramidal** | 73 | -.10 | .008 | .02 | .00 | 1 | 1.0 | .02 | .57 | .81 |

***Table S2.*** *Expression – thinning statistics. Main statistics from the expression – thinning correlation analysis at the age of interest (i.e. where thinning significantly related to cell type-specific expression; at young age (5 – 9 years), middle age (35 -41 years), and old age (63-67 years)). Genes indicate the number of genes included in each cell type group. r indicates the average expression - thinning correlation coefficient across all genes within a cell type group. "p" and "p-fdr" denote the significance of the average expression - thinning correlation - estimated as a function of the empirical null distribution – before and after FDR-adjustment for n = 9 different cell type comparisons. See also* ***Figure 3****.*

| | | | Young | | | Middle age | | | Old | | |
|---|---|---|---|---|---|---|---|---|---|---|---|
| **Subclass** | **Main class** | **Genes(n)** | **r** | **P** | **rank** | **R** | **P** | **rank** | **r** | **P** | **Rank** |
| **Astro1** | Astrocyte | 21 | .26 | **<.001** | 2 | .18 | **<.001** | 2 | -.16 | **.007** | 2 |
| **Astro2** | Astrocyte | 21 | .27 | **<.001** | 1 | .22 | **<.001** | 1 | -.15 | **.01** | 5 |
| **CA1Pyr1** | CA1.Pyramidal | 21 | .01 | .15 | 12 | | | | -.12 | .05 | 7 |
| **CA1Pyr2** | CA1.Pyramidal | 21 | .21 | **.001** | 4 | | | | -.21 | **<.001** | 1 |
| **CA1PyrInt** | CA1.Pyramidal | 21 | .12 | .06 | 9 | | | | -.12 | .05 | 6 |
| **CA2Pyr2** | CA1.Pyramidal | 21 | .06 | .34 | 18 | | | | -.06 | .29 | 14.5 |
| **Mgl1** | Microglia | 21 | .20 | **.002** | 5 | | | | -.15 | **.01** | 4 |
| **Mgl2** | Microglia | 21 | .16 | **.01** | 7 | | | | -.11 | .08 | 8 |
| **Pvm1** | Microglia | 21 | .24 | **<.001** | 3 | | | | -.15 | **.008** | 3 |
| **Pvm2** | Microglia | 21 | .17 | **.008** | 6 | | | | -.08 | .20 | 12 |

***Table S3.*** *Cell type subclasses. Relationship between cortical thinning profiles (at young, middle age and old age periods) and gene expression associated with subclasses of astrocytes, microglia, and CA1 pyramidal cells. Rank indicates the ranking of a given cell type subclass in terms of average expression - thickness correlation coefficients (n = 47 subclasses; ranking in the older period has been reversed). Bold p-values indicate significance after FDR-adjustment for multiple comparisons (n = 10 for young and old and n = 2 for the middle-age thinning profiles). Note that in middle-age, only astrocyte subclasses were tested, as, in this period, microglia and CA1 pyramidal-specific gene expression was not associated with cortical thinning (see* ***Figure 2****).*

| | | **ADNI** | | | **AIBL** | | |
|---|---|---|---|---|---|---|---|
| | nGenes | AD-HC | AD-MCI | MCI-HC | AD-HC | AD-MCI | MCI-HC |
| **Astrocyte** | 54 | **-0,29 (.00)** | **-0,29 (.00)** | **-0,27 (.00)** | **-0,29 (.00)** | **-0,25 (.00)** | **-0,33 (.00)** |
| **CA1.Pyramidal** | 103 | **-0,25(.00)** | **-0,26(.00)** | **-0,23(.00)** | **-0,24(.00)** | **-0,23(.00)** | **-0,24(.00)** |
| **Endothelial** | 57 | 0,05(.56) | 0,04(.65) | 0,05(.49) | 0,01(.84) | 0,02(.82) | 0,01(.87) |
| **Ependymal** | 84 | -0,10(.14) | -0,11(.16) | -0,09(.13) | -0,12(.11) | -0,11(.110) | -0,11(.15) |
| **Interneuron** | 100 | -0,02(.75) | -0,02(.72) | -0,01(.84) | -0,03(.75) | -0,03(.82) | -0,03(.74) |
| **Microglia** | 48 | **-0,27(.00)** | **-0,26(.00)** | **-0,26(.00)** | **-0,24(.00)** | **-0,21(.00)** | **-0,26(.00)** |
| **Mural** | 25 | -0,08(.55) | -0,08(.52) | -0,07(.49) | -0,07(.66) | -0,67(.76) | -0,09(.55) |
| **Oligodendrocyte** | 60 | 0,07(.38) | 0,07(.47) | 0,08(.27) | 0,03(.75) | 0,02(.82) | 0,05(.67) |
| **S1.Pyramidal** | 73 | 0,11(.14) | 0,10(.16) | 0,12(.08) | 0,10(.19) | 0,09(.21) | 0,10(.19) |

**Table S4.** *Virtual Histology in AD. Relationship between cell-specific gene expression profiles and the cortical decline associated with AD clinical diagnosis (AD vs. HC; AD vs. MCI; MCI vs. AD). Bold p-values indicate significance after FDR-adjustment for multiple comparisons (n = 9). AD = Alzheimer's Disease, MCI = Mild Cognitive Impairment, HC = Healthy Control. See also* ***Figure 4****.*

| Cohort/scanner | Observations | Participants | Obs./participant (mean) | Sex (female:male) | Age (SD) | Age range |
|---|---|---|---|---|---|---|
| **Brazil** | 1174 | 809 | 1.5 | 350:459 | 12.0 (2.6) | 6.8 - 18.9 |
| **Brazil 1** | 573 | 391 | 1.5 | 186:205 | 12.1 (2.7) | 7.0 – 18.9 |
| **Brazil 2** | 601 | 418 | 1.4 | 164:254 | 12.0 (2.5) | 6.9 - 18.7 |
| **SPS** | 548 | 548 | 1 | 300:248 | 49.4 (5.0) | 36.4 - 65.4 |
| **AIBL** | 739 | 435 | 1.7 | 246:189 | 72.6 (6.5) | 60.0 - 92.0 |
| **AIBL 1** | 349 | 189 | 1.8 | 104:85 | 74.0 (7.0) | 60.0 – 92.0 |
| **AIBL 1.1** | 108 | 108 | 1 | 64:44 | 72.5 (5.0) | 63.0 – 90.0 |
| **AIBL 2** | 282 | 140 | 2.0 | 79:61 | 71.0 (5.9) | 63.0 – 91.2 |

***Table S5**. Replication MRI sample descriptives. Main sociodemographic information of the MRI sample, also grouped by scanner. Obs. = Observations.*

| Cohort/scanner | Observations | Participants | Obs./participant (mean) | Sex (female:male) | Age (SD) | Age range |
|---|---|---|---|---|---|---|
| **ADNI** | 6680 | 1725 | 3.9 | 774:951 | 75.1(7.2) | 54.4 - 95.6 |
| **ADNI**$_{HC}$ | 2075 | 570 | 3.6 | 291:279 | 75.8(6.2) | 56.0 – 95.6 |
| **ADNI**$_{MCI}$ | 3097 | 931 | 3.3 | 383:548 | 74.3(7.6) | 54.4 – 93.4 |
| **ADNI**$_{AD}$ | 1508 | 586 | 2.6 | 250:336 | 75.8(7.5) | 54.4 – 93.4 |
| **AIBL** | 1069 | 631 | 1.7 | 281:350 | 73.6 (6.8) | 55.0 - 95.7 |
| **AIBL**$_{HC}$ | 785 | 461 | 1.7 | 200:261 | 73.0(6.5) | 59.8 – 91.6 |
| **AIBL**$_{MCI}$ | 149 | 112 | 1.3 | 61:51 | 75.7(6.9) | 60.0 – 95.7 |
| **AIBL**$_{AD}$ | 135 | 94 | 1.4 | 42:52 | 74.9(8.0) | 55.0 – 92.9 |

***Table S6.*** *AD MRI sample descriptives. Main sociodemographic information of the AD MRI sample group by clinical diagnostic. Note that clinical diagnostic of a given individual may change over time. HC = Cognitively Healthy Controls; MCI = Mild Cognitive Impairment; AD = Alzheimer's disease.*
